# Functional Contribution of Mesencephalic Locomotor Region Nuclei to Locomotor Recovery After Spinal Cord Injury

**DOI:** 10.1101/2022.08.22.504420

**Authors:** Marie Roussel, David Lafrance-Zoubga, Nicolas Josset, Maxime Lemieux, Frederic Bretzner

**Author notes:** Corresponding author: Dr. Frederic Bretzner.

## Abstract

Spinal cord injury (SCI) results in a disruption of information between the brain and the spinal locomotor circuit. Although the spinal cord contains all the neural circuits to generate locomotion, people with SCI are unable to walk due to the absence of descending commands from the brain. Electrical stimulation of supraspinal locomotor centers, such as the Mesencephalic Locomotor Region (MLR), can promote locomotor recovery in acute and chronic SCI rodent models. Although clinical trials are currently underway in SCI patients, there is still debate about the organization of this supraspinal locomotor center and which anatomical correlate of the MLR should be targeted to promote functional recovery. Combining kinematics, electromyographic recordings, anatomical analysis, and mouse genetics, our study reveals that glutamatergic neurons of the cuneiform nucleus contribute to locomotor recovery by enhancing motor efficacy in flexor and extensor hindlimb muscles, and by increasing locomotor rhythm and speed on a treadmill, over ground, and during swimming in mice with chronic SCI. In contrast, glutamatergic neurons of the pedunculopontine nucleus slow down locomotion. Therefore, our study identifies the cuneiform nucleus and its glutamatergic neurons as a therapeutical target to improve locomotor recovery in patients living with SCI.

**One Sentence Summary:** Glutamatergic neurons of the mesencephalic locomotor region contribute to spontaneous locomotor recovery following spinal cord injury and selective activation of a discrete glutamatergic subpopulation in this region can further improve functional outcome in chronic spinal cord injury.

## INTRODUCTION

Although the spinal cord contains all the circuitry necessary to locomotion, people with Spinal Cord Injury (SCI) are unable to walk due to the absence of commands from the brain. Motor recovery can be partially achieved by rehabilitative training and neuromodulatory therapies intended to promote the descending motor command from the brain to the spinal cord after SCI (Bonizzato et al., 2018; Capogrosso et al., 2016; Wagner et al., 2018; Wenger et al., 2016; Bonizzato and Martinez, 2021; Bachmann et al., 2013; Bonizzato et al., 2021). Recently, deep brain stimulation of the Mesencephalic Locomotor Region (MLR), a supraspinal locomotor center, has been shown to improve locomotor functions in rats with chronic but incomplete SCI with even a few spared axonal fibers (Bachmann et al., 2013; Bonizzato et al., 2021). Interestingly, these functional changes come with an extensive reorganization in the brainstem region after SCI (Zörner et al., 2014), thus supporting the important contribution of the MLR to locomotor recovery after incomplete SCI.

The anatomical correlate of this functional region has been initially identified as the cuneiform nucleus (CnF), a cluster of glutamatergic neurons; and the pedunculopontine nucleus (PPN), a cluster of glutamatergic and cholinergic neurons. However, there is still debate about the exact anatomical correlate of this supraspinal locomotor center. Previously, using a head-restrained mouse on an air-lifted ball (Lee et al., 2014a; Roseberry et al., 2016), it was shown that optogenetic stimulation of glutamatergic neurons of the MLR (including the CnF and PPN) can generate locomotion in contrast to cholinergic or GABAergic MLR neurons. More recently, using smaller volumes of adeno-associated virus to circumscribe photo-stimulation to a nucleus of interest, it was shown that glutamatergic CnF neurons can initiate locomotion in freely behaving mice (Josset et al., 2018; Caggiano et al., 2018), whereas glutamatergic PPN neurons induce a few unreliable episodes of locomotion only at long latency upon high-frequency stimulation (Caggiano et al., 2018). Furthermore, glutamatergic CnF neurons accelerate locomotor rhythm and speed during ongoing locomotion, whereas glutamatergic and cholinergic PPN neurons only prolong the stance phase, contributing to postural adjustments and slowing locomotor rhythm in normal conditions (Josset et al., 2018). With ongoing clinical trials aiming to assess deep brain stimulation in the vicinity of the PPN of patients with incomplete spinal cord injury (SCI) (Stieglitz LH, 2017) and DBS in the PPN (Bonizzato et al., 2021) or the CnF (Hofer et al., 2022) improving locomotor recovery in animal models of SCI, it is now urgent to gain a better understanding of how these distinct neuronal populations of the midbrain can contribute to and promote functional locomotor recovery after SCI.

We hypothesize here that glutamatergic neurons of the CnF and PPN contribute to spontaneous locomotor recovery after SCI and that selective activation of glutamatergic CnF, but not glutamatergic PPN neurons, can further improve locomotor functions. Combining detailed kinematics, electromyographic (EMG) recordings, anatomical analyses, and mouse genetics, we found that glutamatergic CnF neurons modulate locomotor pattern and rhythm and enhance motor efficacy in ipsilesional hindlimb muscles after SCI, in contrast to glutamatergic neurons of the PPN. As a therapeutical approach, we also found that long trains of photo-stimulation delivered above glutamatergic CnF neurons promote functional recovery of initiation and locomotion after SCI, whereas glutamatergic PPN neurons slow down locomotion, thus identifying the CnF and its glutamatergic neuronal population as a neurosurgical target to promote functional locomotor recovery in patients with SCI.

## RESULTS

### Mice exhibit asymmetrical locomotor pattern following incomplete SCI

Adult mice underwent a lateral hemisection at the low thoracic level, abolishing supraspinal inputs from the brain on the left side of the lumbar spinal cord controlling hindlimb locomotion. Using kinematic and electromyographic recordings, we assessed spontaneous locomotor recovery after SCI (Fig. 1, Fig. S1 and Table S1). As previously reported in other animal species (Barrière et al., 2008; Barrière et al., 2010; Martinez et al., 2012b; Martinez et al., 2012a; Courtine et al., 2008; Helgren and Goldberger, 1993; Kuhtz-Buschbeck et al., 1996), mice initially displayed a transient paralysis on the side of the SCI over the first week post-SCI (Fig. 1C-D and Fig. 1G-H) with a decrease in toe movement and forward foot placement (Fig. 1E-F) and in the motor activity of the flexor, which were co-activated with otherwise weak extensors (Fig. 1G-J). This likely contributed to hind-paw dragging during swing and a very short stance. Although this limb displayed some locomotor-like movements, there was loss of weight support and no plantar stepping.

**Fig. 1:**
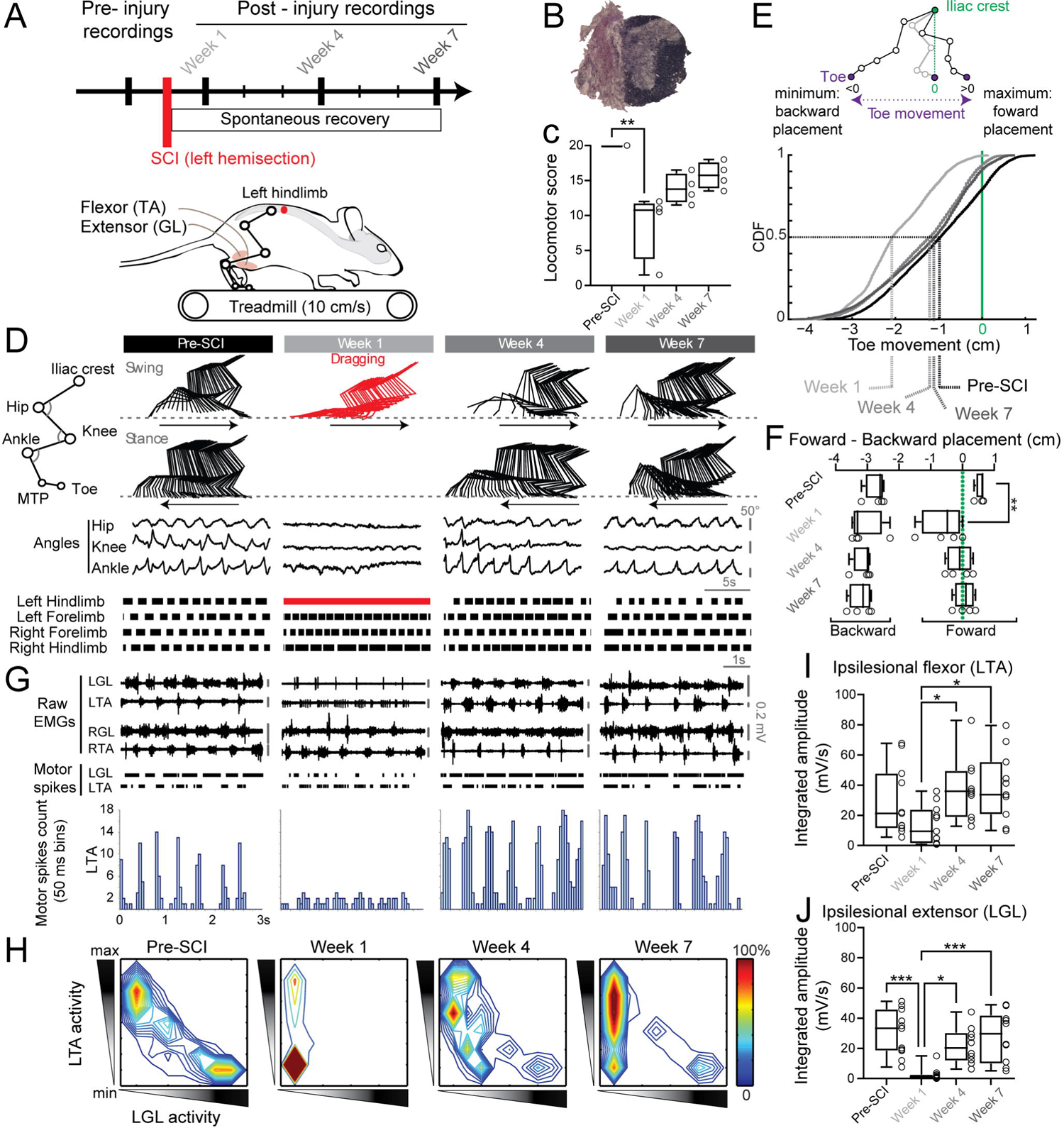
Spontaneous locomotor recovery after spinal cord injury (SCI). (A) Kinematics and electromyographic (EMG) recordings during treadmill locomotion before and after SCI. (B) Example of a hemisection at the left thoracic level (T8-T9) with cresyl violet and luxol Fast Blue staining. (C) Locomotor score of the ipsilesional (left) hindlimb before and after SCI (n=4 mice). (D) Stick diagrams of the ipsilesional hindlimb during the swing and stance phase (arrows indicate movement direction), joint angles, and gait diagrams (bars represent the stance). (E) Cumulative distribution function (CDF) of paw movement in reference to the iliac crest before and after SCI (n=4 mice). (F) Forward and backward paw placement in reference to the iliac crest (n=4 mice). (G) Raw EMGs and raster of motor spikes of flexor and extensor muscles. LTA: left tibialis anterior, LGL: left gastrocnemius lateralis, RTA: right tibialis anterior, RGL: right gastrocnemius lateralis. (H) Flexor activity as a function of extensor activity of the ipsilesional hindlimb (n=9 mice). Note the loss of alternation between flexor and extensor 1 week after SCI and its recovery associated with a longer flexor activity at 4 and 7 weeks after SCI. (I) Integrated amplitude of the ipsilesional left flexor (n=11 mice). (J) Integrated amplitude of the ipsilesional left extensor (n=11 mice). *P < 0.05, **P < 0.01, ***P < 0.001.

Eventually, within a few weeks after SCI, the ipsilesional limb (i.e., on the lesion side) increased its stance duration and extensor activity, in addition to exhibiting better coordination in its flexor (Fig. 1I-J). This is turn reduced hind-paw dragging during swing and improved plantar stepping ability (Fig. 1D and 1G), thus contributing to locomotor recovery of the ipsilesional limb over time. These kinematic and motor changes were accompanied by changes in the contralesional limb (i.e., opposite to the lesion side). Indeed, the contralesional limb modified its locomotor pattern accordingly by reducing its flexor activity and swing duration while increasing its extensor activity and stance duration (Fig. 1D-G, Fig. S1). These kinematic and motor changes persisted up to 7 weeks after SCI, contributing to postural adjustments on the contralesional hindlimb and functional stepping recovery of the ipsilesional hindlimb.

### No anatomical reorganization of medullar projecting MLR neurons after SCI

Given that the medullary reticular formation relays MLR inputs during locomotion (Noga et al., 1991; Noga et al., 2003; Shefchyk et al., 1984) and that SCI leads to an extensive reorganization of projections between midbrain and brainstem nuclei after SCI (Zörner et al., 2014), we hypothesized an asymmetrical reorganization of the connectivity between glutamatergic neurons of MLR nuclei and their postsynaptic medullary targets following incomplete SCI. To test this hypothesis, a retrograde tracer Fast Blue was injected stereotaxically in the contralesional medullary reticular formation of adult transgenic mice 7 weeks after SCI or sham surgery (Fig. 2A; mice with Fast Blue injections leaking in the ipsilesional brainstem were excluded from our analysis, Fig. S2). We specifically targeted the Gigantocellular Reticular Nucleus, the alpha and ventral portion of the Gigantocellular Reticular Nucleus, and the Lateral Paragigantocellular Nucleus, which are important to the motor command (Bretzner and Brownstone, 2013; Lemieux and Bretzner, 2019; Bouvier et al., 2015; Capelli et al., 2017). Combining immunohistochemistry and stereological techniques (Fig. 2b), we identified and quantified the neurotransmitter phenotype (e.g., glutamatergic, cholinergic, or both glutamatergic/cholinergic) of back-labeled neurons in ipsi- and contralesional CnF and PPN nuclei.

**Fig. 2:**
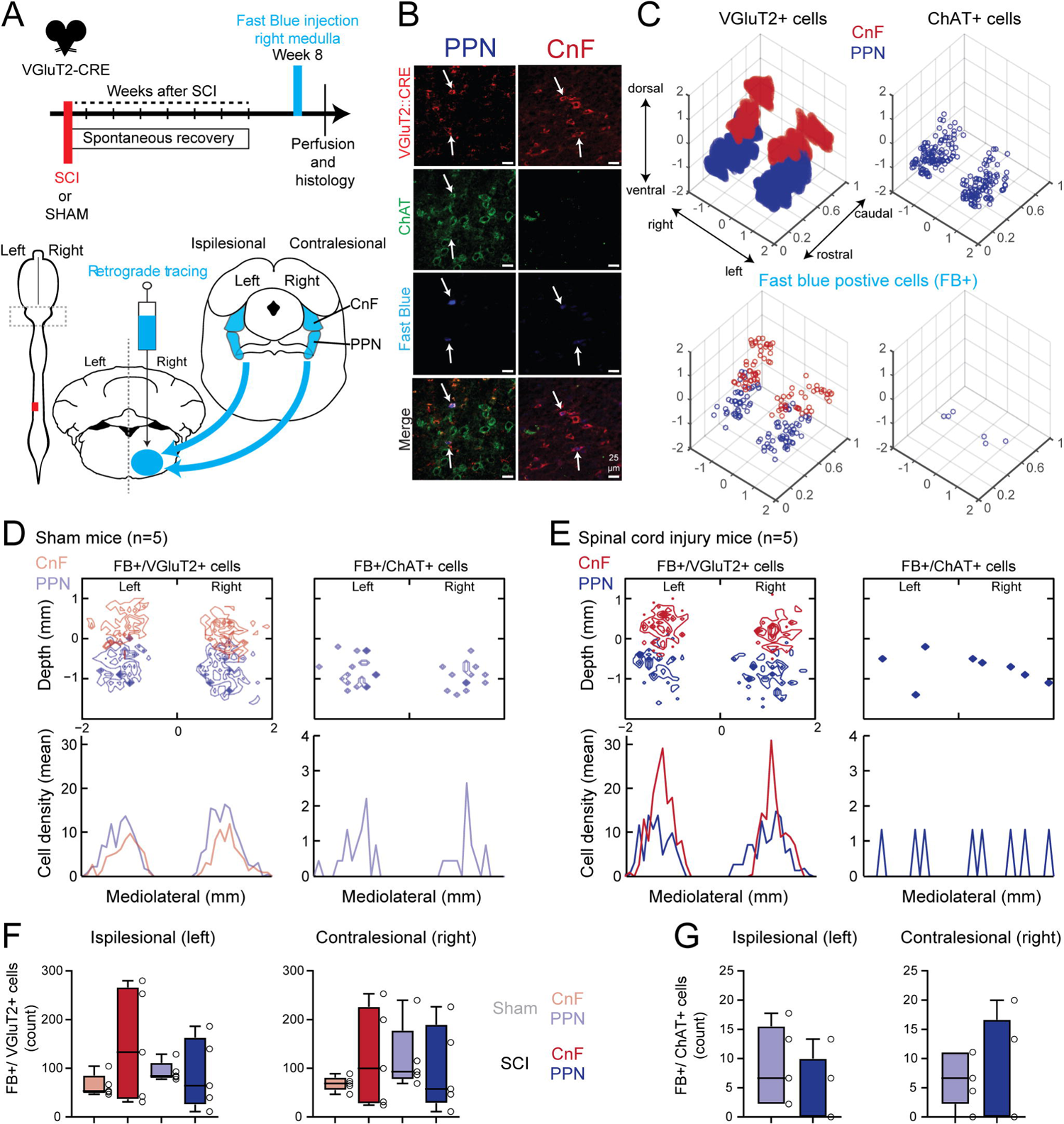
No anatomical change in the number of glutamatergic and cholinergic CnF and PPN neurons projecting to the motor medulla after chronic SCI. (A) Injection of a retrograde tracer, Fast Blue, in the contralesional medulla 8 weeks after SCI or after laminectomy without SCI (n=5 SCI and 5 SHAM mice). (B) Examples of immunostaining of glutamatergic (VGluT2::CRE in red) and cholinergic (ChAT in green) neurons with retrograde labelling (Fast Blue in blue) in the cuneiform nucleus (CnF) and pedunculopontine nucleus (PPN). (C) Example of a spatial representation of glutamatergic neurons (VGluT2+), cholinergic neurons (ChAT+), and back-labeled glutamatergic (VGluT2+/FB+) and cholinergic (ChAT+/FB+) neurons in the ipsi- and contralesional CnF and PPN of a chronic mouse. (D-E) 2D mediolateral and dorsoventral distributions (top) and mediolateral projection (bottom) of back-labeled glutamatergic and cholinergic mesencephalic neurons from SHAM (D) and SCI (E) mice. (F-G) Mean and SD of cell counts of glutamatergic (F) or cholinergic (G) retrogradely labelled neurons in the CnF and PPN of SHAM and SCI mice.

Using the most ventral part of the 4^th^ ventricle as a reference, our 3D and 2D reconstructions illustrate a bilateral and symmetrical organization of CnF and PPN nuclei according to their neurotransmitter phenotype (e.g., glutamatergic, cholinergic, or double) and their unilateral projection in the contralesional medullary reticular formation (Fig. 2C-E). We found a high density of medullary projecting glutamatergic neurons in both left and right CnF and PPN, but very few cholinergic cells within the PPN (Fig. 2C-E), suggesting a bilateral organization of brainstem-projecting MLR populations in both sham and SCI mice. As previously reported (Lavoie and Parent, 1994; Luquin et al., 2018; Wang and Morales, 2009; Steinkellner et al., 2019), we also found a few double glutamatergic/cholinergic neurons in the PPN (Fig. S2). Nevertheless, in contrast to a previous SCI study (Zörner et al., 2014), the cell density profile and count of glutamatergic and cholinergic neurons of CnF or PPN nuclei of SCI animals were not significantly different from those of sham mice (Fig. 2D-E), suggesting that the organization of glutamatergic and cholinergic MLR neurons projecting to brainstem locomotor circuits was maintained after SCI.

### Genetic deletion of glutamatergic neurons of the contralesional CnF or PPN impairs spontaneous locomotor recovery in chronic SCI mice

Given the functional role of the MLR to locomotor recovery after SCI (Zörner et al., 2014; Bachmann et al., 2013), we next evaluated the requirement of glutamatergic neurons of specific MLR nuclei to functional recovery (Fig. 3, Fig. S3, Fig. S4 and Table S1). To test this requirement, locomotor functions were assessed before and after conditional genetic ablation of glutamatergic neurons of the contralesional CnF or PPN in chronic mice that had spontaneously recovered locomotor functions 8 weeks after SCI (Fig. 3A). We ensured that both CnF and PPN groups exhibited a similar behavioral recovery 8 weeks after SCI (Brown and Martinez, 2018) prior to their genetic ablation, and that both groups showed the same extent of SCI (Fig. 3B). During treadmill locomotion (Fig. 3C-E), genetic ablation of glutamatergic CnF neurons decreased the angular excursion of the ipsilesional hip in 60% of mice (Fig. 3C-D) but did not impair their intralimb and interlimb coordination overall (Fig. 3E and Fig. S3). In contrast, genetic ablation of glutamatergic PPN neurons had almost no significant locomotor effects, except in the coupling between the hip and the ankle (Fig. 3E).

**Fig. 3:**
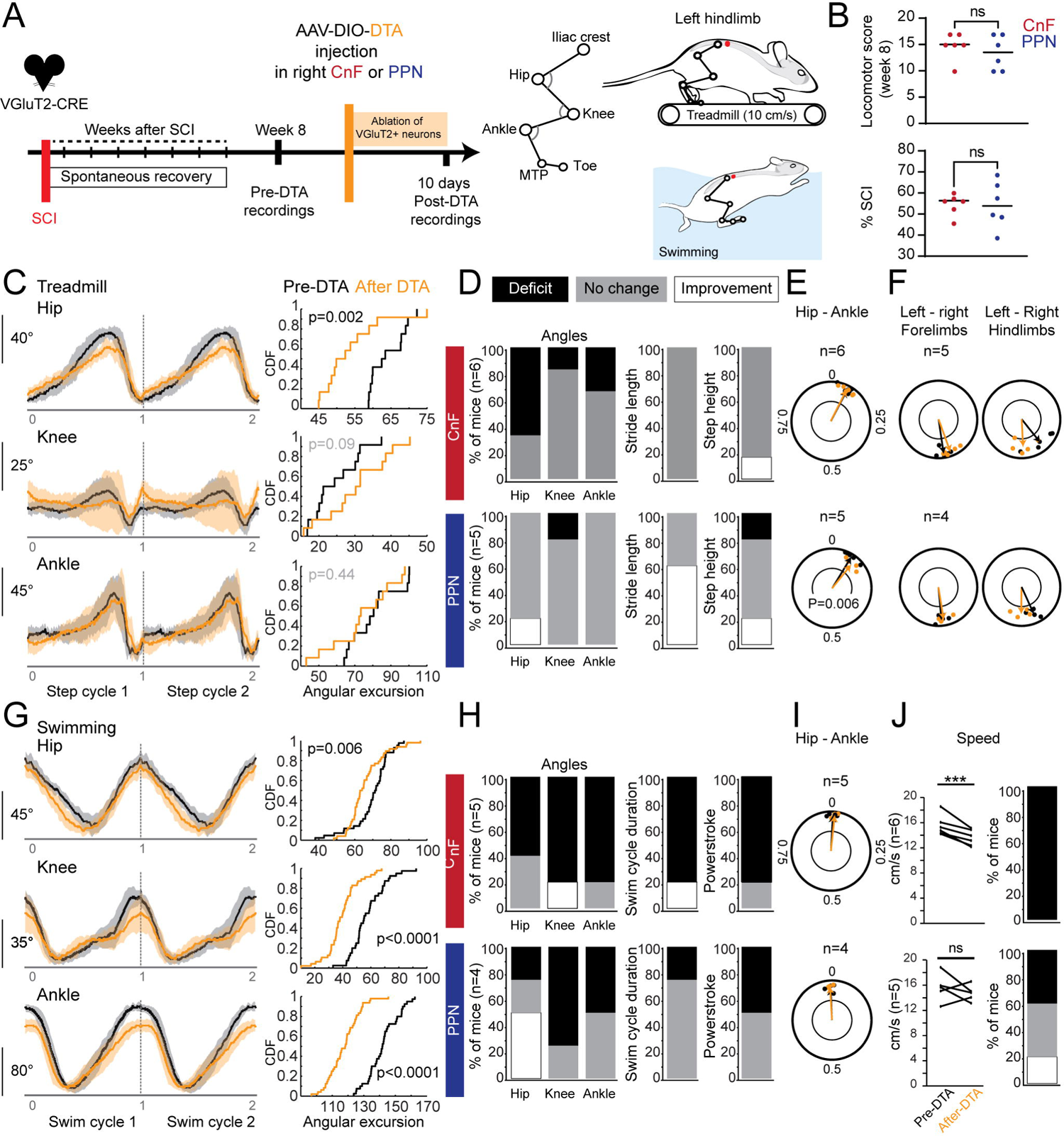
Genetic ablation of contralesional glutamatergic CnF or PPN neurons impairs spontaneous locomotor recovery after chronic SCI. (A) Kinematic analysis before and after genetic ablation of glutamatergic neurons of the contralesional right CnF or PPN in chronic SCI mice. (B) Locomotor score and extent of the lesion site in both experimental groups. (C) Mean and SD of hip, knee, and ankle joint angles of the ipsilesional hindlimb during treadmill locomotion before and after genetic ablation of glutamatergic CnF neurons in a mouse. Cumulative density frequency (CDF) of the angular excursion of hindlimb joints. (D) Percentage of mice with significant decrease (deficit), increase (improvement), or absence of change in the angular excursion of the joints, stride length, and step height. (E) Coupling between the angle of the ipsilesional hip and ankle. (F) Bilateral fore- and hindlimb coupling (anchored on the right forelimb and hindlimb, respectively). (G) Mean and SD of hip, knee, and ankle joint angles of the ipsilesional hindlimb during swimming before and after genetic ablation of glutamatergic CnF neurons in a mouse. CDF of the angular excursion. (H) Percentage of mice with significant deficit, improvement, or absence of change in the angular excursion, swim cycle duration, and power stroke. (I) Coupling between the angle of the ipsilesional hip and ankle. (J) Average speed before and after genetic ablation and associated percentage of mice with deficit. ***P < 0.001.

During swimming (Fig. 3G-I), however, genetic ablation of glutamatergic neurons significantly decreased the angular excursion of the hip, knee, and ankle of the ipsilesional hindlimb in 60 to 80% of chronic CnF mice and in 20 to 80% of chronic PPN mice (Fig. 3H). Furthermore, genetic deletion of glutamatergic CnF neurons decreased the step cycle duration and power stroke in 80% of CnF mice (Fig. 3G) and swimming speed in all CnF mice (Fig. 3I), whereas fewer mice were impaired upon genetic ablation of glutamatergic PPN neurons. These findings argue that glutamatergic neurons of the contralesional CnF are more important than their PPN counterparts to spontaneous locomotor recovery after SCI.

### Glutamatergic CnF neurons contribute to spontaneous locomotor recovery after SCI

Having shown that genetic ablation of glutamatergic neurons of the contralesional CnF or PPN is necessary to spontaneous locomotor recovery after SCI, we next assessed their functional contribution by evaluating changes in motor efficacy throughout the step cycle (Fig. 4A). The extent of lesion size and changes over time in locomotor score were similar for both CnF and PPN groups (Brown and Martinez, 2018) (Fig. 4B and Fig. S5).

**Fig. 4:**
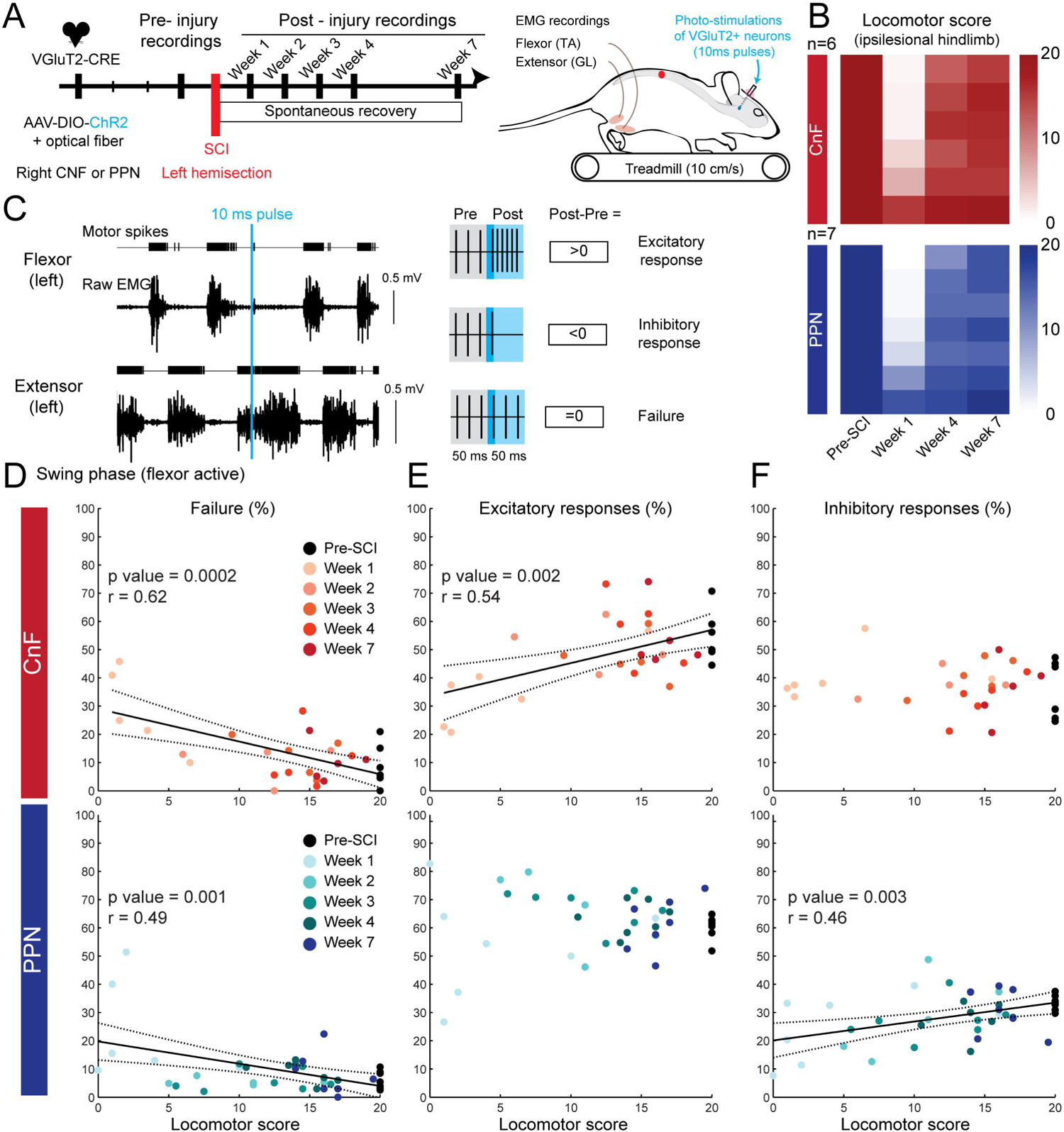
Glutamatergic neurons of the CnF contribute more efficiently than those of the PPN to spontaneous locomotor recovery of the ipsilesional hindlimb after SCI. (A) Combination of electromyographic recordings with short pulses (10ms) of photo-stimulation to probe changes in motor efficacy of glutamatergic CnF or PPN neurons to locomotor recovery after SCI. (B) Color-coded matrices illustrating the locomotor score of the ipsilesional left hindlimb of mice before and after SCI. (C) Example of background electromyographic (EMG) activities of the ipsilesional flexor (tibialis anterior) and extensor (gastrocnemius lateralis) muscles during treadmill locomotion with a 10ms pulse photo-stimulation. The difference in the density of motor spikes evoked post versus pre photo-stimulation identified whether responses were excitatory, inhibitory, or absent (indicating a failure). (D-F) Proportion of failure (D), excitatory (E), and inhibitory (F) motor responses evoked in the ipsilesional flexor muscle during the swing phase as a function of the locomotor score of the ipsilesional hindlimb over the course of spontaneous locomotor recovery after SCI.

To assess changes in motor efficacy, the percentage of failure, excitatory, and inhibitory phase-dependent EMG responses were measured in the ipsilesional flexor and extensor muscles over a 50 ms time window upon photo-stimulation of 10 ms pulse duration delivered during locomotion at steady and comfortable speed before and after SCI (Fig. 4C). An increase in the number of motor spikes indicated excitatory motor responses, whereas a decrease indicated inhibitory motor responses and an absence of change indicated a failure. We quantified changes over time in the proportion of motor responses in the flexor muscle during the swing phase (Fig. 4) and in the extensor during the stance phase (Fig. S6) as function of the locomotor score of the ipsilesional hindlimb. There was a high failure rate in motor responses in both flexor and extensor muscles 1 week after SCI (Fig. 4D and Fig. S6A), which returned eventually toward pre-injury levels over time while the animals resumed spontaneous locomotor recovery. Indeed, changes in the failure rate in both muscles correlated negatively with the locomotor score of the ipsilesional hindlimb (Fig. 4D and Fig. S6A): the lower the failure rate, the higher the locomotor score, thus suggesting a transient interruption of the descending motor drive after SCI.

Regarding the flexor muscle during the swing phase (Fig. 4E-F), changes in the proportion of excitatory motor responses correlated significantly only with changes in locomotor score upon photo-stimulation of glutamatergic CnF neurons. In contrast, changes in the proportion of inhibitory motor responses correlated only with changes in the locomotor score upon photo-stimulation of glutamatergic PPN neurons, suggesting that glutamatergic CnF neurons by their excitatory drive contribute more efficiently than the PPN to motor recovery of the flexor muscle during the swing phase after SCI.

Regarding the extensor muscle during the stance phase (Fig. S6B and C), correlations between changes in the proportion of inhibitory motor responses and locomotor scores were positive upon activation of either the CnF or PPN (Fig. S6C). However, changes in the proportion of excitatory responses correlated significantly and positively with changes in the locomotor score upon photo-stimulation of glutamatergic CnF neurons, whereas the correlation was negative upon photo-stimulation of glutamatergic PPN neurons (Fig. S6B), thus supporting overall a higher excitatory efficiency of the CnF over the PPN in spontaneous recovery of the extensor muscle during the stance phase after SCI.

Overall, changes in the density of motor spikes in excitatory and inhibitory motor responses were positively correlated with changes in locomotor score in both flexor and extensor muscles upon photo-stimulation of the CnF or PPN (Fig. S7 and Fig. S9). However, changes in the amplitude of motor spikes were only correlated with the locomotor score in excitatory responses of the flexor evoked upon glutamatergic CnF neurons (Fig. S8). Taken together, these changes in the excitatory motor drive and behavior suggest that glutamatergic CnF neurons contribute more efficiently than PPN neurons to spontaneous recovery of stepping ability after SCI.

### Glutamatergic neurons of the CnF promote initiation of locomotion in chronic SCI mice

Having shown that glutamatergic neurons of both the CnF and PPN are necessary and contribute to some extent to spontaneous recovery after SCI, we next investigated whether stimulation of one of these neuronal populations would be more efficient in promoting initiation of locomotion after chronic SCI (Fig. 5, Fig. S11, Fig. S12 and Table S1). As illustrated by their trajectory in open field (Fig. 5A), long trains of photo-stimulation delivered above glutamatergic CnF neurons generated consistent episodes of locomotion (Fig. 5B and 5D: 100% of trials at 20 Hz and 84% at 50 Hz) with long distance displacement (Fig. 5A and 6D). As illustrated by body direction (Fig. 5C), the first 500 ms of stimulation generated already straight locomotion (Fig. 5C) at very short latency (Fig. 5E). Although long trains of photo-stimulation delivered above glutamatergic PPN neurons generated bouts of locomotion, these episodes were often inconsistent and unreliable (Fig. 5I, 5K and 5J: 23% of trials at 20 Hz and 34% at 50Hz) and always occurred with a long latency after the end of the photo-stimulation (Fig. 5L), as previously reported (Caggiano et al., 2018). Regarding body direction, whereas glutamatergic CnF neurons generated straight locomotion with a displacement over the first 500 ms of photo-stimulation (Fig. 5c and 6f: 100% of trials at 20 Hz and 84% at 50Hz), glutamatergic PPN neurons in contrast failed and more systematically evoked head rotation without body displacement as illustrated by shorter arrows (Fig. 5J and 5M: 3% of trials at 20 Hz and 5% at 50Hz). In summary, sole activation of glutamatergic neurons of the CnF generates straight locomotion in chronic SCI mice.

**Fig. 5:**
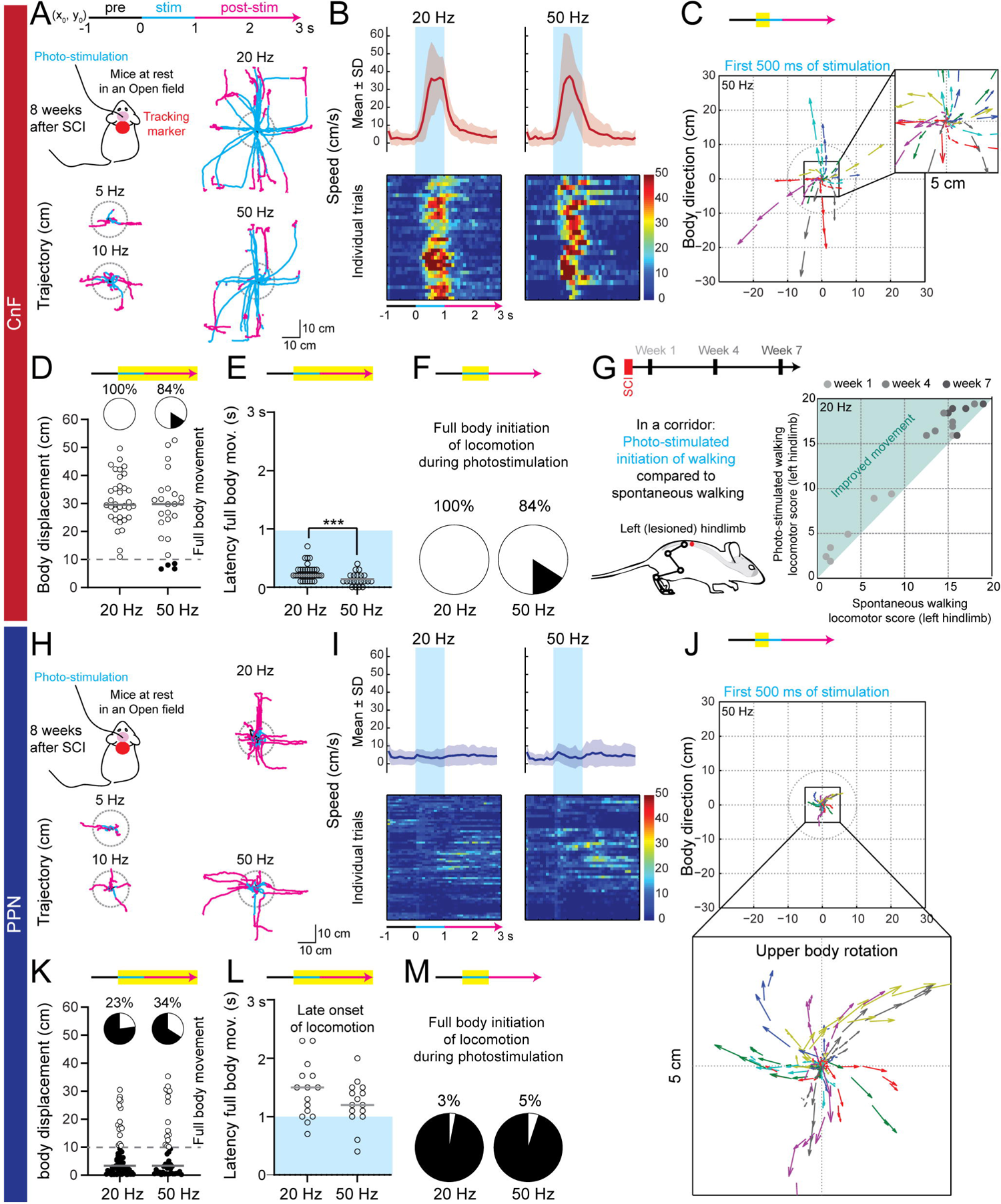
Initiation of locomotion evoked by stimulation of glutamateric CnF neurons after chronic SCI. (A) Full body movement trajectories evoked upon long-train photo-stimulation of glutamatergic neurons of the CnF at different frequencies (n = 5 mice). Mice were tracked using a marker placed on their neck. Trajectories of all mice were centered at (x0,y0) 1 s prior to stimulation. Gray circles represent a 10 cm radius, distance from which a full body initiation was considered. (B) Mean and SD of locomotor speed evoked upon photo-stimulation. Color-coded matrices representing individual trials (100 ms bins). (C) Vectors represent the upper body direction and the distance travelled (vector length) during the first 500 ms of photo-stimulation at 50 Hz (yellow highlight on the timeline). (D) Body displacement (and median) evoked by photo-stimulation. A 10 cm displacement was considered as a full body movement (i.e., an initiation of locomotion). All displacements occurring within the 3 seconds from the start of the stimulation were considered. (E) Latency to initiate full body movement upon photo-stimulation. (F) Percentage of trials in which initiation of locomotion started within the 1st second of photo-stimulation. (G) Ipsilesional hindlimb locomotor score during spontaneous versus evoked by CnF photo-stimulation (20Hz) walking over time after SCI (n = 7). (H) Trajectories of full body movement evoked upon photo-stimulation of glutamatergic neurons of the PPN (n = 7 mice). (I) Mean and SD of locomotor speed. (J) Vectors illustrate consistent upper body rotation ipsilateral to the stimulation site within a 5 cm radius. 50 Hz photo-stimulation of the PPN evoked consistent rapid motor movements with little displacement, considered as a postural rotation. (K) Body displacement. (L) Latency to initiate full body movement started after the end of the PPN photo-stimulation. (M) Percentage of trials in which initiation of locomotion started within the 1st second of photo-stimulation of the PPN. ***P < 0.001.

**Fig. 6:**
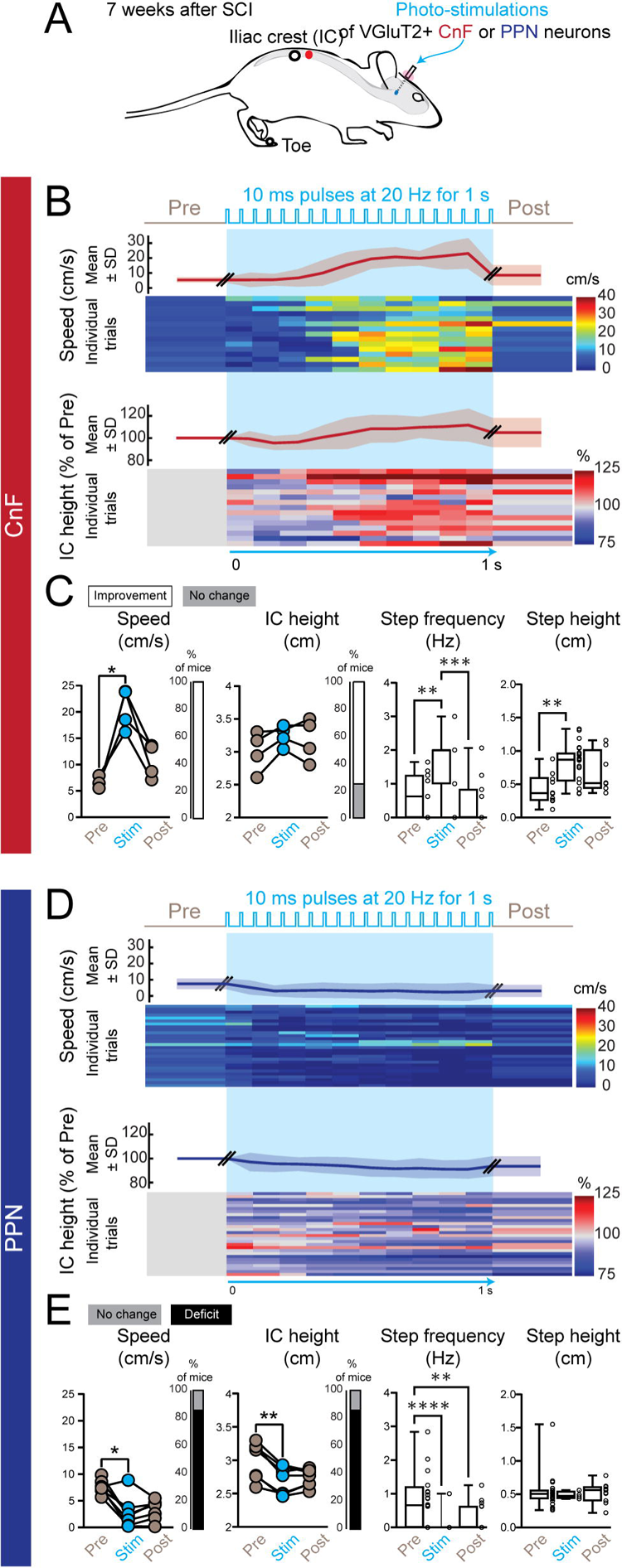
Functional improvement during overground locomotion upon long trains of photo-stimulation of glutamatergic CnF neurons after chronic SCI. (A) Trains of photo-stimulation (10ms pulses at 20 Hz for 1s) delivered in glutamatergic neurons of the CnF (n = 4) or PPN (n = 7) 8 weeks after SCI. (B) Time course of locomotor speed (mean and SD) and individual trials (below, 3-4 trials per mouse, 100 ms bins). Time course of the Iliac crest height (mean and SD) and individual trials normalized on data prior to photo-stimulation (below). (C) Mean of locomotor speed of individual mouse evoked upon photo-stimulation of glutamatergic CnF neurons (500ms pre/stim/post periods, last 500ms of the stimulation episode). Percentage of mice exhibiting a significant increase in speed upon photo-stimulation in comparison to the pre-stimulation period. Mean height of the iliac crest of each mouse. Percentage of mice exhibiting a significant increase in the height of the iliac crest upon photo-stimulation. Step frequency and step height of all step cycles combined. (D) same as B. (E) Mean of locomotor speed of individual mouse evoked upon photo-stimulation of glutamatergic PPN neurons. Percentage of mice exhibiting a deceleration upon photo-stimulation of the PPN. Mean of the height of the iliac crest. Percentage of mice exhibiting a decrease in the height of the iliac crest upon photo-stimulation of the PPN. Step frequency and step height of residual steps occurring upon photo-stimulation of the PPN. **P < 0.01, ***P < 0.001, ****P < 0.0001.

### Activation of glutamatergic CnF neurons improves posture and recovery of basic and voluntary stepping in chronic SCI mice

Having shown that glutamatergic neurons of the CnF initiate episodes of locomotion in animals at rest, in contrast to the PPN (Fig. 5), we next investigated how activation of these neurons can modulate posture and voluntary locomotion in a corridor. As shown by the height of the iliac crest to the toe (Fig. 6), glutamatergic neurons of the CnF increased the posture, step height, and step frequency, and enhanced the locomotor speed of chronic SCI mice during ongoing locomotion, whereas neurons of the PPN decreased locomotion and postural tone (Table S1). Similar observations were also reported during treadmill locomotion at a comfortable steady speed (Fig. 7B). Moreover, changes in postural tone increased linearly as a function of changes in locomotor speed: photo-stimulation of glutamatergic CnF neurons increased speed and posture, whereas glutamatergic PPN neurons reduced these parameters (Fig. S13), thus suggesting that these two neuronal populations could modulate gait and posture.

**Fig. 7:**
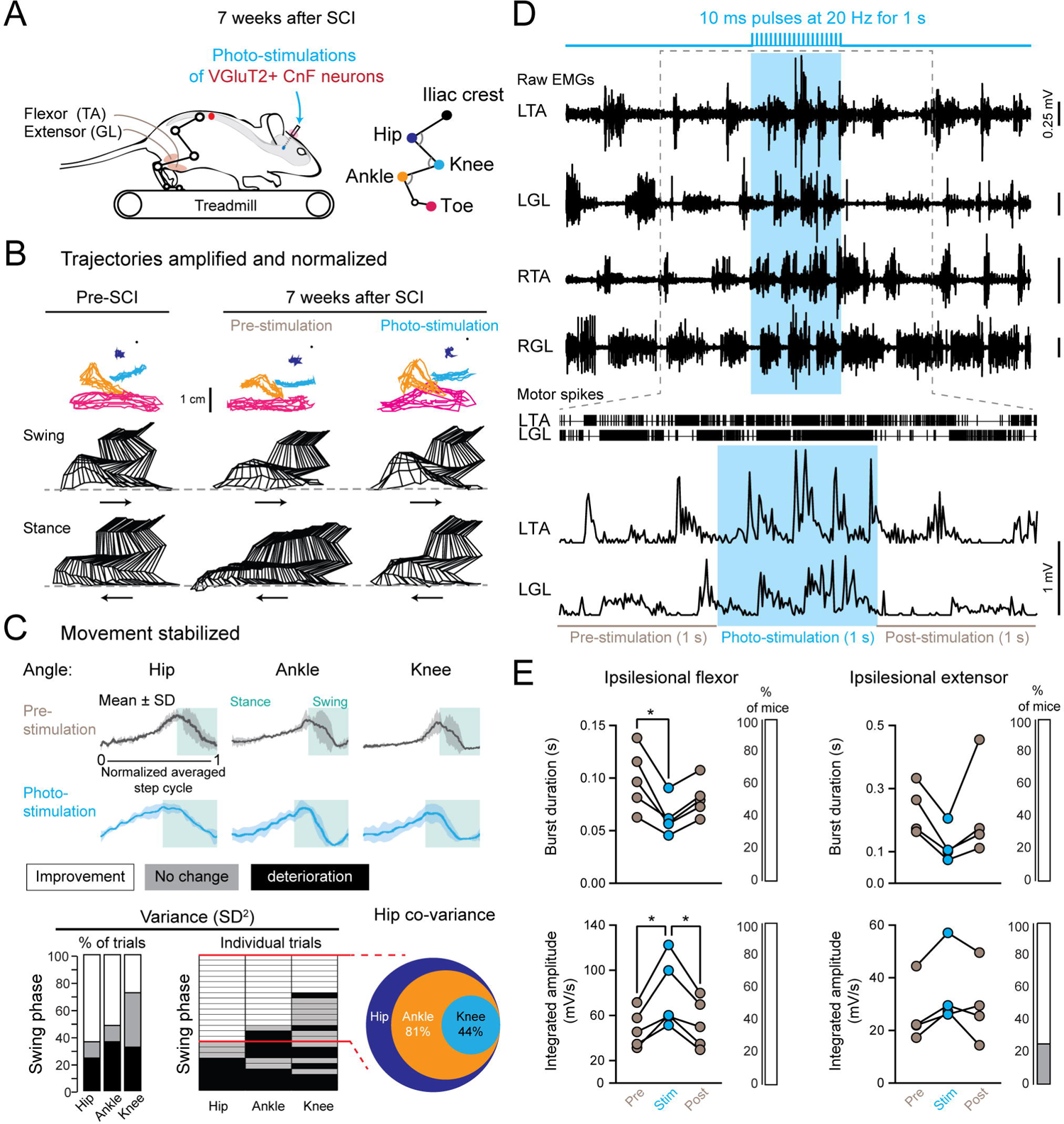
Functional improvement of intralimb coordination during treadmill locomotion upon long trains of photo-stimulation of glutamatergic CnF neurons after chronic SCI. (A) Kinematic and EMG recordings of the ipsilesional hindlimb evoked upon long trains of photo-stimulation (10 ms at 20 Hz for 1 s) of glutamatergic CnF neurons during treadmill locomotion 7 weeks after SCI (n = 5 mice). (B) Cyclic trajectories of hindlimb joints (anchored on the iliac crest) during locomotion before and after chronic SCI. Stick diagrams illustrate the swing and stance phases (arrows show the direction of movement). (C) Mean and SD of the hip, knee, and ankle joint angles during a normalized step cycle. Green areas indicate the swing phase of the step cycle. Percentage of step cycles in which photo-stimulation induced a decrease (improvement) or increase (deterioration) in the variance of hip, knee, and ankle joint angles during the swing phase (left). Individual trials were aligned according to the evolution of the variance of the hip angle (middle). Pie chart showing that ankle and knee stabilization occurred only when the hip was stabilized (right). (D) Raw EMG recordings (top), raster of motor spikes of the LTA and LGL (middle), and rectified EMG traces of the LTA and LGL at higher magnification (below). (E) Burst duration and integrated amplitude of background EMG bursts in the ipsilesional flexor and extensor. Percentage of mice showing a significant improvement (white) or no change (gray). *P<0.05. Abbreviations: see Fig. 1 legend.

SCI patients exhibit higher variability in their intralimb coordination that impedes their complete recovery (Easthope et al., 2018; Sohn et al., 2018). We therefore hypothesize that by decreasing this variability, photo-stimulation of the glutamatergic neurons of the CnF might improve stepping ability. To further investigate that possibility, we analyzed kinematic changes of the intralimb coordination during treadmill locomotion at steady and comfortable speed of chronic SCI mice (Fig. 7). While the standard deviation of the hip, knee, and ankle joints was especially high during the swing phase of locomotion prior to stimulation (black traces in Fig. 7C), activation of the glutamatergic CnF neurons significantly decreased this variability (blue traces in Fig. 7C) with the strongest changes in the hip and ankle followed by the knee (Fig. 7C, 60% of trials in the hip, 50% in the ankle, and 25% in the knee). Interestingly, this decreased variability during the swing phase co-varied according to the variance of the hip. Indeed, when the variance of the hip was significantly reduced, the variability of the ankle joint decreased significantly in 81% of trials and that of the knee in 44% of trials, thus supporting smoother and steadier stepping. Furthermore, analyses of electromyographic recordings also showed that activation of glutamatergic neurons of the CnF decreased the burst duration of flexor and extensor muscles of both ipsi- and contralesional hindlimbs (Fig. 7D-E for the ipsilesional hindlimb and Fig. S14 for the contralesional hindlimb, Table S1), thus contributing to an increased locomotor speed. Stimulation also increased the burst amplitude of the ankle dorsiflexor activity (e.g., Tibialis Anterior), contributing to increasing the step height and toe clearance during the swing phase of locomotion. Taken together, these results suggest that stimulation of glutamatergic neurons of the CnF can efficiently modulate the spatio-temporal recruitment of muscles, smoothen and stabilize the intralimb joint coordination of the ipsilesional hindlimb, and enhance speed, thus improving overall locomotor recovery after chronic SCI.

### Unloading the body weight improves locomotor functions driven by glutamatergic CnF neurons in chronic SCI mice

Although sensory feedback participates to functional locomotor recovery after SCI, unloading the body weight (even partially) is often combined with physical training to promote motor recovery in incomplete SCI patients and animal models (Alcobendas-Maestro et al., 2012; de Leon and Dy, 2017). Therefore, we tested whether long trains of photo-stimulation delivered above glutamatergic neurons of either the contralesional CnF or PPN can promote locomotor function while unloading the animal’s weight during swimming in chronic SCI (Fig. 8; Fig. S15, Table S1).

**Fig. 8:**
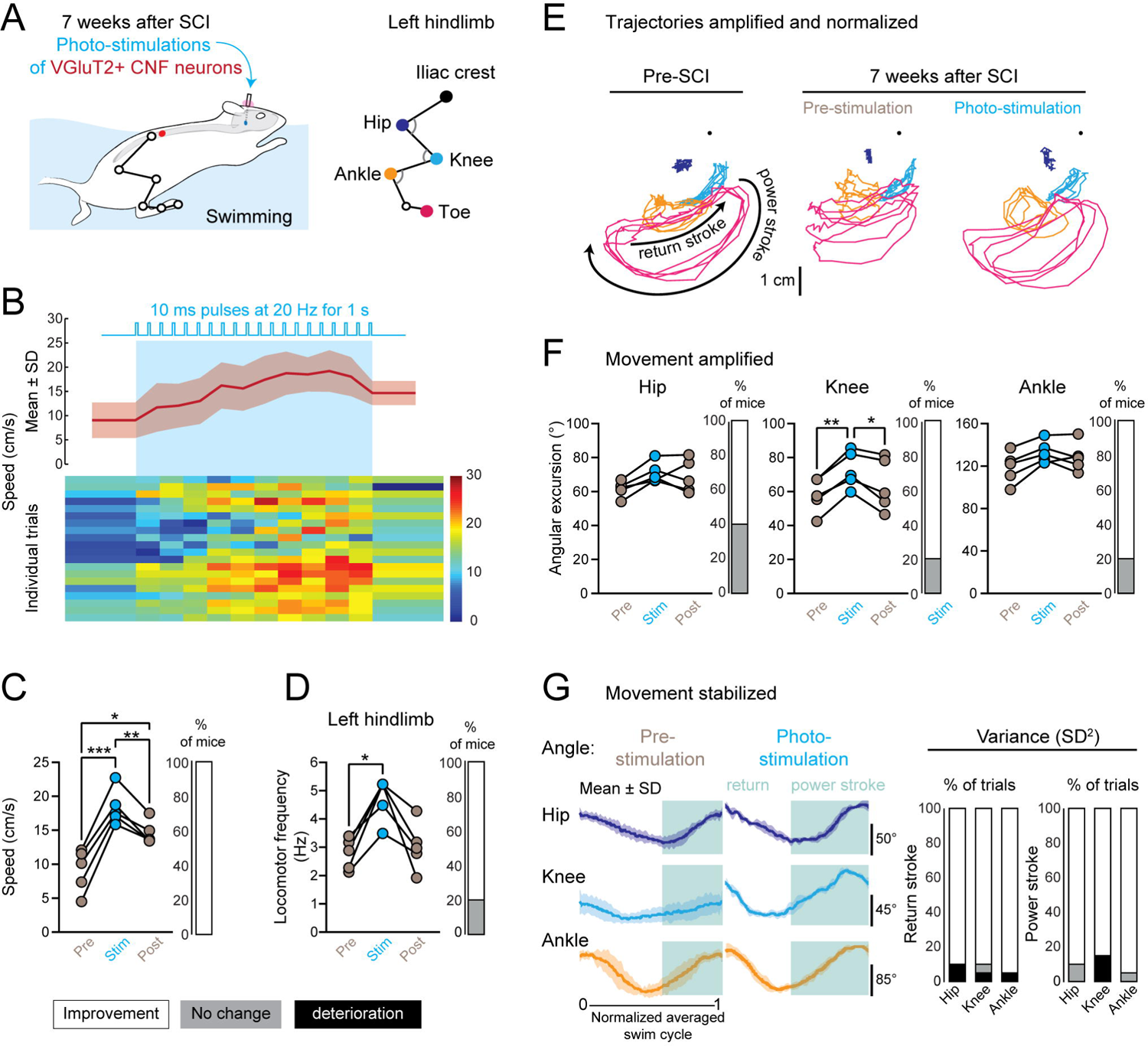
Functional improvement of swimming upon long trains of photo-stimulation of glutamatergic CnF neurons after chronic SCI. (A) Kinematic analysis of the ipsilesional hindlimb evoked upon long trains of photo-stimulation (10 ms at 20 Hz for 1s) of glutamatergic CnF neurons during swimming 7 weeks after SCI (n = 5 mice). (B) Mean and SD of the swimming and individual trials color-coded as a function of speed (100 ms bins, 4 trials per mouse). (C) Mean swimming speed (500ms pre/stim/post periods, last 500ms of the 1s stimulation). 100% of mice exhibited increased speed upon photo-stimulation. (D) Mean swimming frequency upon photo-stimulation. Percentage of mice exhibiting increased locomotor frequency upon photo-stimulation. (E) Trajectories of the hip, knee, ankle, and toe anchored to the iliac crest during a swimming bout before and 7 weeks after SCI. (F) Mean angular excursion of the ipsilesional hip, knee, and ankle. Percentage of mice exhibiting a significant increase in their angular excursion upon photo-stimulation (60% for the hip and 80% for the knee and ankle). (G) Mean and SD of joint angles during normalized swim cycles upon photo-stimulation. Percentage of trials exhibiting a significant decrease in the variance of joint angles during return and power strokes upon photo-stimulation. *P < 0.05, **P < 0.01, ***P < 0.001. (H)

Seven weeks after SCI, activation of glutamatergic CnF neurons increased swimming speed and frequency (Fig. 8A-D) and improved the trajectory of the hindlimb joints (Fig. 8E), in addition to increasing the angular excursion of the hip in 60% of mice and of the knee and ankle in 80% of mice (Fig. 8F). Furthermore, activation of the CnF also improved fluidity in movement during the power stroke and return stroke of the ipsilesional hindlimb in more than 90% of trials in the hip, knee, and ankle joints (Fig. 8G). In contrast, activation of glutamatergic neurons of the PPN decreased swimming speed and had very little effect on intralimb coordination of the ipsilesional hindlimb (Fig. S15). Taken together, these results show that activation of glutamatergic CnF neurons promote more efficient and smoother intralimb coordination of the ipsilesional hindlimb, thus improving functional locomotor recovery of the ipsilesional hindlimb of chronic SCI mice.

## DISCUSSION

Our results reveal new concepts that fundamentally alter the understanding of the distinct contribution of MLR nuclei to functional locomotor recovery following SCI and their potential as deep brain stimulation targets to promote rehabilitation in patients with chronic SCI: (I) despite an absence of neuroanatomical changes within the brainstem, (ii) both glutamatergic neurons of the CnF and PPN contribute functionally differently to locomotor recovery; (iii) glutamatergic neurons of the CnF are more important than those of the PPN for spontaneous locomotor recovery; (iv) if glutamatergic PPN neurons contribute to the functional inhibitory drive in both flexor and extensor muscles of the ipsilesional hindlimb, only glutamatergic neurons of the CnF contribute to the functional excitatory drive of both flexor and extensor muscles; moreover (v) activation of glutamatergic neuronal populations of the CnF with long trains of photo-stimulation at 20 Hz improve the intralimb coordination of the ipsilesional hindlimb, as well as gait and posture during basic locomotion while stepping on a treadmill and during voluntary locomotion while walking on the ground or swimming in a pool in chronic SCI. Overall, our results argue that clinical trials should consider targeting the CnF rather than the PPN, and especially glutamatergic neurons of the CnF, in patients with SCI.

### The PPN is a complex structure that is suboptimal for generating locomotion

Using animal models, the PPN has been initially identified as an anatomical correlate of the mesencephalic locomotor region based on electrical stimulation and postmortem reconstructions (Garcia-Rill et al., 1985; Skinner et al., 1990; Garcia-Rill and Skinner, 1987). Given that Parkinson’s disease (PD) patients typically display difficulties in initiating and executing locomotor movements, with rigidity, tremor, and postural instability (Morris et al., 2001; Morris et al., 1996), deep brain stimulation (DBS) of the PPN has been investigated in advanced Parkinsonian patients with motor complications who are refractory to pharmacological treatments. Although PPN stimulation improves gait and postural adjustments in some Parkinsonian patients (Mazzone et al., 2005; Plaha and Gill, 2005), the functional outcomes have been extremely variables across clinical studies (Thevathasan et al., 2018). Such variability, also reported in several animal models of Parkinson’s (Goetz et al., 2016; Rauch et al., 2010), has recently raised questions about the efficacy of the stimulation parameters and the anatomical correlates of the MLR.

Although there is still no consensus regarding the most appropriate stimulation parameters, as little is known about the neural mechanisms activated by DBS in the vicinity of the PPN (Lozano et al., 2019; Garcia-Rill et al., 2019), recent optogenetic studies of glutamatergic neurons of the PPN have shown functional discrepancies in animal models with locomotor initiation, deceleration, head rotation, and even anxious-like behaviors (Caggiano et al., 2018; Josset et al., 2018; Carvalho et al., 2020; Kroeger et al., 2017), that could reflect the complexity of this nucleus. Indeed, in addition to the multiple pathways running through it, the PPN exhibits an extreme divergence in its presynaptic inputs and postsynaptic projections (Martinez-Gonzalez et al., 2011; Caggiano et al., 2018). As shown by recent optogenetic studies, neuronal populations within the PPN can exhibit distinct functional effects according to their postsynaptic projection. If little is known about medullary-projecting PPN neurons, which will be the population most likely involved in locomotion, there is evidence that substantia nigra-projecting PPN neurons generate grooming and handling (Ferreira-Pinto et al., 2021), striatal-projecting PPN axons generate head rotation (Assous et al., 2019), and spinally projecting PPN neurons induce locomotor arrest and rearing (Ferreira-Pinto et al., 2021), thus arguing that the PPN is a complex neurological structure with a wide range of functional motor outcomes.

### Glutamatergic neurons of the CnF enable stepping ability after SCI without discomfort

Recently, electrical stimulation of the PPN has been shown to initiate bipedal locomotion of chronic rats contused at the thoracic level (Bonizzato et al., 2021). However, electrical PPN stimulation also appears to induce pain as reported by a grimace test in these chronic SCI animals. Similarly, using optogenetic tools in the mouse, we found that photoactivation of glutamatergic neurons of the PPN could also induce some discomfort in chronic SCI mice, but not to the extent of SCI rats upon electrical stimulation of the PPN. Although we cannot exclude some discrepancies between both animal species and SCI models, electrical stimulation is not as specific as optogenetic activation in recruiting different neuronal populations, axons of passage, and unwanted neural structures in the vicinity of the electrode. By their caudal location to the PPN, electrical stimulation of the PPN might have recruited the kolliker/parabrachial nuclei involved in grimaces and aversive responses during pain (Raver et al., 2020; Roeder et al., 2016). In contrast, optogenetic stimulation of glutamatergic neurons, especially in the CnF, resulted in almost no or very little apparent discomfort in chronic SCI mice.

Electrical stimulation of the CnF has also been recently shown to promote quadrupedal locomotion of chronic thoracic SCI rats (Hofer et al., 2022). Using a less severe thoracic SCI, we also found that optogenetic photo-stimulation of glutamatergic neurons of the CnF with long trains of 20 Hz for 1 s increased postural tone and initiated locomotion in chronic SCI mice, with functionally better stepping ability and smoothness in locomotion than during spontaneous episode of locomotion. Although stimulation of glutamatergic neurons of the PPN usually failed to evoke locomotion with our standard stimulation protocol at 20Hz, increasing the stimulation frequency up to 50 Hz induced very slow locomotor movements. These, however, were not reliable and occurred only at very long latency after the end of the stimulation before and after SCI, as previously shown in uninjured mice (Caggiano et al., 2018). Overall, our findings suggest that the CnF and especially glutamatergic neurons of the CnF are a better target than the PPN for promoting functional recovery of locomotor initiation in chronic SCI.

### Glutamatergic CnF neurons are necessary and sufficient to improve functional recovery during locomotion

Genetic anterograde and retrograde tracing studies have shown a direct connection between glutamatergic neurons of both the CnF and PPN nuclei and the medulla (Ferreira-Pinto et al., 2021; Caggiano et al., 2018). Pharmacological, lesion, cooling, and genetic deletion studies have previously shown that reticulospinal pathways of the medullary reticular formation relay supraspinal MLR inputs to the spinal locomotor circuit (Shefchyk et al., 1984; Noga et al., 1991; Noga et al., 2003; Capelli et al., 2017). Not surprisingly, reticulospinal pathways appear to contribute to functional locomotor recovery following SCI (Filli et al., 2014; Zörner et al., 2014; Engmann et al., 2020; Asboth et al., 2018), presumably by relaying MLR and/or cortical inputs. Although anatomical reorganization has been reported between medullary-projecting midbrain neurons following cervical SCI (Zörner et al., 2014) and axonal midbrain sprouting in the medulla after thoracic SCI (Hofer et al., 2022), we found no anatomical changes in genetically identified glutamatergic medullary-projecting CnF or PPN nuclei after thoracic SCI, despite functional behavioral and motor improvements. Indeed, genetic deletion of glutamatergic neurons of the CnF or PPN impaired spontaneous locomotor recovery during walking and swimming, but deficits were stronger upon genetic ablation of CnF neurons than those of the PPN. Similarly, motor efficacy measurements also revealed that excitatory motor responses evoked in ipsilesional flexor and extensor muscles by glutamatergic neurons of the CnF correlated robustly with changes in locomotor recovery. This was not the case with the PPN, suggesting that glutamatergic neurons of the CnF contribute more efficiently than those of the PPN to spontaneous recovery of locomotor functions following SCI.

### Glutamatergic CnF neurons promote functional locomotor recovery in chronic SCI mice

As previously shown before SCI (Josset et al., 2018), activation of glutamatergic CnF neurons increased postural tone as well as locomotor pattern and rhythm in chronic SCI mice during treadmill locomotion, as well as during voluntary locomotion while walking in a corridor or swimming in a pool, whereas activation of glutamatergic PPN neurons usually slowed locomotor rhythm, eventually inducing locomotor arrests. Interestingly, similar decelerations and stops have also been recently reported upon photo-stimulation of PPN neurons projecting to the spinal cord in intact animals (Ferreira-Pinto et al., 2021).

Despite very little understanding of the neural brainstem networks underlying locomotion (Ausborn et al., 2019; Roussel et al., 2020), recent optogenetic studies have shown that photo-stimulation of glutamatergic neurons of the LPGi modulate locomotor rhythm (Capelli et al., 2017), presumably by relaying glutamatergic inputs of the CnF, whereas photoactivation of glutamatergic or glutamatergic V2a (Lhx3/Chx10) expressing neurons of the Gi reset locomotor rhythm and induce locomotor arrests (Bouvier et al., 2015; Cregg et al., 2020; Lemieux and Bretzner, 2019; Usseglio et al., 2020), presumably by relaying glutamatergic inputs of the PPN. Further studies are needed to genetically dissect the neural mechanisms of these medullary nuclei to motor recovery after SCI. Overall, our results support the hypothesis that glutamatergic CnF neurons are more efficient than those of the PPN in improving stepping ability of the ipsilesional hindlimb, in addition to enhancing gait and posture, in chronic SCI mice during stereotyped and voluntary locomotion.

## Conclusion

With a current clinical trial assessing MLR DBS in patients with incomplete SCI (Stieglitz LH, 2017), our findings reveal that the CnF offers a better and more reliable neurological target in comparison to the PPN for promoting recovery of motor and locomotor functions, strengthening the importance of evaluating the role of the CnF and especially glutamatergic neurons of the CnF in patients suffering from SCI. Our results also highlight the importance of continuing our genetic dissection of functional microcircuits within PPN in biomedical research. Although optogenetic and optical technologies are still in their infancy regarding first-in-human clinical trials targeting more accessible neural structures to restore sensory loss (Sahel et al., 2021; Dieter et al., 2020), new technological advances might soon allow better control of deep brain neuronal populations to promote recovery of gait and posture in humans suffering from SCI or neurodegenerative diseases, such as Parkinson’s disease or Amyotrophic Lateral Sclerosis. In summary, our current findings in an animal model of SCI suggest that DBS of the CnF or optogenetic activation of glutamatergic CnF neurons should be further investigated in chronic SCI patients.

## MATERIALS AND METHODS

### Mice

VGluT2-IRES-Cre (RRID: IMSR_JAX:016963) mouse strain was maintained on a mixed genetic background (129/C57Bl6). Given the prevalence and severity of spinal cord injury in human and rodent males (Lee et al., 2014b) and the weaker functional recovery in comparison to female rodents (Datto et al., 2015), adult (≥60 days) male mice weighing approximately 30 g were used in this study. Before experiments, mice were housed in groups with a maximum of 5 per cage. Mice with brain implants were housed individually after surgery to avoid implant damage. AAV2/9 EF1-DIO-hChR2(H134R)-mCherry (titer 9E12 GC/ml to 1.2E13 CC/ml) or AAV2/9-EF1a-mCherry-flex-dTA (titer 1.2E13 CC/ml) was injected in VGluT2-IRES-Cre mice to induce a restricted cre-lox recombination (Canadian Neurophotonics Platform Viral Vector Core Facility (RRID:SCR_016477)). Housing, surgery, behavioral experiments, and euthanasia were performed in compliance with the guidelines of the Canadian Council on Animal Care and approved by the local committee of Université Laval (CPAUL3).

### AAV injections and optical fiber implantations

Under isoflurane (1,5%–2% O2) anesthesia, the mouse was installed in a stereotaxic frame; a craniotomy was performed for AAV injection and chronic implantation of a unilateral optical fiber (diameter: 200 μm) above the nucleus of interest in photostimulated groups. The targets were the cuneiform nucleus (CnF; anteroposterior from the Bregma (AP), −4.6 mm; mediolateral (ML), 1.25 mm; depth, 2.2 mm) or the pedunculopontine nucleus (PPN; AP, −4.6 mm; ML, 1.25 mm; depth, 3.25 mm). Chronically implanted groups underwent the AAV injection (100 nL) and the subsequent implantation of an optical fiber in the same surgery. A glass micropipette (WPI, ID: 0.53 and OD: 1.19 mm) was backfilled with mineral oil and fixed on a nano-injector (Nanoliter 2010 Injector, WPI). The pipette was lowered slowly into the nucleus of interest. After a 5-min period, the AAV was injected at a rate of 50nL/min. To avoid any leakage of the AAV, the glass pipette was held in place for 5 min following the injection before being slowly retracted. The optical fiber (200 µm) was held in place with dental acrylic (cat# 525000 and 526000, A-M Systems) and machine screw (cat#19010-10, FST, North Vancouver, Canada).

### Spinal cord injury

Under isoflurane (1.5%–2% O2) anesthesia, the mouse was installed in a stereotaxic frame; a laminectomy was performed with or without (SHAM group) spinal lesion. The enlargement of vertebra T7 was used as a landmark and T7 was removed using a Friedman-Pearson rongeur to access the T8-T9 spinal segments. In spinalized animals, the left spinal cord was transected dorso-ventrally (the medio-lateral landmark was the posterior spinal vein) using a 30G needle. To prevent regeneration, a piece of absorbable cellulose gauze hemostat was inserted in the lesion site. Axial muscles and skin were replaced and sutured. For all surgical procedures, analgesics (Buprenorphine hydrochloride SR: 5mg/kg) were provided at the end of the surgery for long duration release. After a 1-week recovery, mice were tested in the laboratory.

### Retrograde tracing

8 weeks after SCI, under isoflurane (1,5%–2% O2) anesthesia, the mouse was installed in a stereotaxic frame; a craniotomy was performed for retrograde tracer injection: Fast Blue (EMS-Chemie, Gross-Umstadt, Germany, 50 nL, 2% suspension in phosphate buffer with 2% dimethyl sulfoxide) in the contralesional medulla: AP = 5.5 mm; L = 0.5 mm; D = 5.5 mm. A 2 µL Hamilton neuro syringe was used to perform this injection. To avoid leakage, the neuro syringe was kept in place for 2 min and removed slowly after the injection. Mice were perfused 4 days after the Fast Blue injection.

### Locomotor tests

Mice were tested during the day in a room dedicated to behavioral experiments. Before any surgery, mice were trained to swim and walk on a treadmill. During swimming, treadmill, and spontaneous walking, reflective markers were placed on each joint of the left ipsilesional hindlimb (iliac crest, hip, knee, ankle, and toe). Mice were filmed on both sides with high-frequency cameras (Genie HM640, Dalsa Teledyne; 250 frames/s). In the open field test, mice were filmed from above (100 frames/s) and a reflective marker was placed on their neck to track body movements. Videos were digitized with StreamPix 6.0 (Norpix) and analyzed offline using custom-designed software, OpenPose (Cao et al., 2021), and MATLAB.

Mice learned to swim in straight lines and reach a platform at the end of a transparent corridor (length: 53 cm, width: 5 cm, height: 18 cm). To prevent hypothermia, the temperature of the water was monitored and maintained between 23 and 25 °C. Three to 4 laps were analyzed per mouse (with or without photo-stimulation).

We tested spontaneous overground walking in a transparent corridor (length: 60, width: 5 cm, height: 25) connecting two open boxes—one transparent and one painted black. Mice were placed in the brighter box and crossed the corridor to find refuge in the darker box. Three to 4 passages were analyzed per mouse (with or without photo-stimulation).

Kinematics and EMGs recordings were performed during two separate sessions on a treadmill (Exer 3/6 treadmill, Columbus instruments). During EMG experiments, mice were filmed from both sides at 100 frames/s without any markers on the hindlimb. Five photo-stimulations were analyzed per mouse. During open field experiments (a rectangular treadmill surface (38 x 43 cm), mice were videotaped from above. A locomotor score was used to monitor spontaneous motor recovery after lateral thoracic hemisection (Brown and Martinez, 2018). Hindlimbs were scored individually each week after SCI. The score on a 20-point scale takes into consideration the articular coordination, the weight support, the digit position, the foot placement while stepping, the tail position, and the fore-hindlimb coordination.

### Optogenetic and electrophysiological recordings

Before and after SCI and throughout all locomotor tests, optical manipulations of Channelrhodopsin-2.0 (ChR2)-expressing neurons were undertaken in freely behaving mice. The pattern and timing of optical manipulations were controlled using a mechanical shutter (Connectorized Mechanical Shutter Adapters; Doric, Canada) and controller (SR470 Laser Shutter Controller; Stanford Research Systems, California, USA) synchronized online during kinematic and EMG recordings. Channelrhodopsin-2.0-expressing neurons (ChR2) were photostimulated by using a blue laser (50 mW power, 473nm wavelength, Laserglow Technologies,Ontario, Canada). Before SCI, a threshold was determined to reliably evoke behavioral responses in each mouse for each locomotor test in 3 separate sessions of photo-stimulation,1 week apart each. During EMG recordings, the laser power threshold was determined using motor responses evoked upon 10 ms pulse photo-stimulations delivered to the animal at rest. Pulses were delivered during treadmill locomotion at the defined threshold (between 1 to 18 mW) at steady locomotor speed (10 cm/s). Regarding the timing of stimulations, continuous 10 ms pulses were delivered every 3 s to assess changes in locomotor pattern (total of 150 pulses); trains of 10 ms pulses at 5-10-20-50 Hz were also used for a duration of 1 s every 5 s.

### EMG activity

EMG activity of the tibialis anterior (TA, ankle flexor), gastrocnemius lateralis (GL, ankle extensor) muscles were recorded using acute electrodes. Mice were anaesthetized with isoflurane (1.5-2%) to insert dual core wires into muscles of interest, as previously described elsewhere (Josset et al., 2018; Lemieux and Bretzner, 2019). Recordings were made when mice were fully awake and ready to walk. EMG signals were amplified (x 1000), band-pass filtered (0.1–10 kHz), sampled at 10 KHz, and digitally converted (Power 1401; CED, Cambridge, UK) using Spike2 version 8 (CED, Cambridge, UK). EMG signals were high-pass filtered, rectified, and analyzed offline using custom-designed software and MATLAB.

### Kinematic analysis

As previously described (Josset et al., 2018; Lemieux and Bretzner, 2019), joint markers of the iliac crest, hip, ankle, and toe were detected. To avoid skin slippage, the knee was inferred by triangulation using the length of the femur and the tibia. Hindlimb joints were detected using the machine learning software OpenPose (Cao et al., 2021). All OpenPose predictions were manually verified and corrected if needed on custom-designed software (Lemieux et al., 2016). Custom MATLAB scripts were used to analyze joints kinematics. Using custom-made software, the beginning of the stance (contact on the ground) and the beginning of the swing (lift from the ground) phases of the hindlimbs during treadmill locomotion were manually tagged. Regarding swimming, the swim cycle was divided into a power and return stroke which were defined using the evolution of the hip angle during swimming. The power stroke ranges from the minimum to the maximum of the hip angle; the return stroke ranges from the maximum to the minimum angle.

### Analysis of background motor activities and motor responses

Motor spikes were extracted from raw EMG recordings using a threshold of 5 times the mean of the background signal at rest. Then, a custom-made MATLAB script was used to identify the beginnings and endings of step cycles using time-sensitive changes in motor spike density. For each hindlimb, the reference muscle was the tibialis anterior. The step cycle was divided in 5 equal epochs, the first two corresponding to the active phase of the flexor (associated with the swing phase of the limb) and the last three corresponding to the active phase of the extensor (associated with the stance phase of the limb). Changes in motor spike density and mean amplitude were assessed as the difference between a 50 ms time window before and after 10 ms photo-stimulation. For the analysis of trains of photo-stimulation or unstimulated locomotion, the burst of activity of the muscles (and the step cycles) were selected manually in Spike2.

### Perfusion

Mice were deeply anesthetized and transcardially perfused with 10 mL saline (0,9% NaCl) followed by 20 mL paraformaldehyde (4% PFA). Tissues (spinal cord and brain) were harvested and post-fixed overnight in 4% PFA, then in 30% sucrose until saturation. Tissues (spinal cords and brains) were frozen in Leica tissue freezing medium, then cut on a Leica cryostat (20 µm slices, Leica CM1860, Germany).

### Immunochemistry

Immunostainings were performed on brainstem slices to confirm the position of the optical probe, the extent of the cre-lox recombination, and analyzed back-labelled neurons from tracing experiments. The following primary antibodies were used: anti-choline acetyltransferase (ChAT) 1:100 (Chemicon-Millipore, AB144P), anti-Cre recombinase (CRE) 1:1000 (EMD Millipore, MAB3120), and the anti-NeuN 1:500 (Chemicon-Millipore, ABN78). The following secondary antibodies were used: donkey anti-mouse-AF594 1:1000 (Thermofisher Scientific, A-21203), donkey anti-goat-AF488 1:1,000 (Abcam, AB150129) donkey anti-rabbit-AF594 (Invitrogen, A21207), and donkey anti-rabbit-AF350 (Invitrogen, A10039).

To confirm optical probe location and cre-lox recombination extent, images were taken on an Axio Imager M2 microscope connected to an AxioCam camera using ZEN2 software (Zeiss, Germany). Low-magnification reconstructions were generated to delineate the extent of the cre-lox recombination and determine the stereotaxic coordinates of the tip of the optical cannula according to anatomical landmarks (superior cerebellar peduncle, inferior colliculus, and the periaqueductal gray) and anatomical atlases (Franklin and Paxinos, 2008; VanderHorst and Ulfhake, 2006). Cholinergic immunostaining was used to identify and localize the cholinergic PPN. The cholinergic staining and the extent of cre-lox recombination were evaluated by outlining the area on low-magnification reconstructions to determine whether the stimulation site was located within the CnF or PPN. Spinal cord lesions were imaged using brightfield microscopy. Using Fiji (Schindelin et al., 2012), the extents of the spinal cord lesion were evaluated on three subsequent slices at the epicenter of the lesion. The area of spared tissue was compared to adjacent intact sections, the extent of the lesion ranged from 40% to 65%.

### Stereology

For tracing experiments, animals in which Fast Blue injection was circumscribed to the right medulla including the gigantocellular reticular nucleus (Gi), the lateral para-gigantocellular reticular nucleus (LPGi), and the ventral pars of the gigantocellular reticular nucleus (GiV) we included for analysis. Every two slices were imaged using an epifluorescence microscope (Olympus BX51, Tokyo, Japan) associated with stereology software (Stereo Investigator, MicroBrightField Bioscience, Colchester, VT). Using anatomical landmarks, the borders of the CnF and PPN were traced. A random subsampling was used, and neurons were counted in selected sections (50 to 150 µm^2^ squares). Sizes of the sections were determined by the surface on the slices and the volume of the nucleus. Taking in consideration the number of counted cells, the interval between sections, the area of the nucleus, and the thickness of the slices, the number of counted cells was estimated. In some experiments, all neurons were counted manually every other section enabling a 3D representation of the PPN and CnF (Fig. 2-c). Custom MATLAB scripts enabled pooling all mice together and representing the spatial distribution (medio-lateral and dorso-ventral) of retrogradely labelled neurons.

### Quantification and statistical analysis

Information about mice numbers, statistical tests, and data representation can be found in Fig. legends. Data are represented as mean ± standard deviation and statistical difference was indicated by asterisks (* *P*≤0.05, ** *P*<0.01, *** *P*<0.001, **** *P*<0.0001). Before every analysis, the normality of the data distribution was assessed using a Shapiro-Wilk test. To test statistical difference from a specified value, we used a one-sample t-test if the distribution was normal or a Mann-Whitney test if the distribution was not normal. To compare groups, a one-way ANOVA was performed followed by a Bartlett post-test if the distribution was normal. Otherwise, if the distribution was not normal, a Kruskall-Wallis test was performed with a Dunn’s multiple comparison post-test. To compare paired groups, we used repeated measures one-way ANOVA followed by Tukey’s comparison test for normally distributed values and a Friedman test followed by Dunn’s multiple comparisons post hoc test otherwise. Boxplots represent the 25^th^ percentile, the median, the 75^th^ percentile, and the whiskers the maximum and minimum values.

## List of Supplementary Materials

Fig. S1: Background EMG activity in the contralesional flexor and extensor muscles after SCI.

Fig. S2: Cell counts of glutamatergic, cholinergic, and glutamatergic/cholinergic CnF and PPN neurons after chronic SCI.

Fig. S3: Intralimb coordination during locomotion upon genetic ablation.

Fig. S4: Genetic ablation of glutamatergic mesencephalic neurons.

Fig. S5: Extent of the cre-lox recombination, site of optical cannulas, and extent of the lesion site.

Fig. S6: Changes in the proportion of failure, excitatory, and inhibitory motor responses in the ipsilesional extensor muscle during the stance phase after SCI.

Fig. S7: Changes in motor spike density of excitatory motor responses evoked in the ipsilesional hindlimb before and after SCI.

Fig. S8: Changes in motor spike amplitude of excitatory motor responses evoked in the ipsilesional hindlimb before and after SCI.

Fig. S9: Motor spike density of inhibitory motor responses evoked in the ipsilesional hindlimb before and after SCI.

Fig. S10: Motor spike amplitude of inhibitory motor responses evoked in the ipsilesional hindlimb before and after SCI.

Fig. S11: No initiation of locomotion upon photo-stimulation at 5 or 10 Hz after chronic SCI.

Fig. S12: No reliable signs of stress or pain upon photo-stimulation of glutamatergic neurons of the CnF or PPN after chronic SCI.

Fig. S13: Speed and posture upon long trains of photo-stimulation of glutamatergic CnF versus PPN neurons after chronic SCI.

Fig. S14: Background EMG activity of the contralesional hindlimb in response to long trains of photo-stimulation of glutamatergic CnF neurons after chronic SCI.

Fig. S15: Long trains of photo-stimulation of glutamatergic PPN neurons failed to improve swimming after chronic SCI.

Table S1: Statistical analysis and p-values.

## Acknowledgments

We would like to thank L. Bihoues, A. Caussaint, G. Gélinas, H. Godet, V. Langevin, S. Harvey, and N. Karimi for their participation in animal handling, data acquisition, and analysis; Dr. J.-F. Lalonde, C. Panter, and P. Drapeau for technical assistance; and M.-C. Richer and S. Bernard for supporting our animal care.

## Author contributions

F.B. conceived and designed the research; M.R., D.L.Z., N.J., and M.L. performed research, and analyzed data; M.R. and F.B. drafted the manuscript; M.R., M.L., and F.B. edited and revised the manuscript; and all authors approved the final version of the manuscript.

## Data and materials availability

All data and custom code associated with this study are present in the paper or the Supplementary Materials will be available upon request.

