## Supplementary material for "Functional Contribution of Mesencephalic Locomotor Region Nuclei to Locomotor Recovery After Spinal Cord Injury"

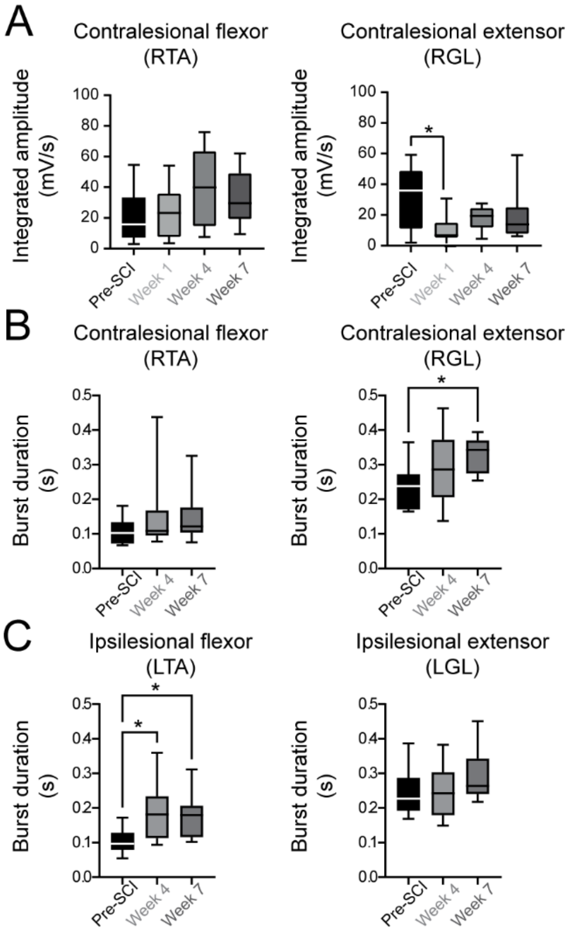


**Fig. S1: Background EMG activity in the contralesional flexor and extensor muscles after SCI.**

(A) Integrated amplitude of the contralesional right flexor and extensor (n=11 mice, Kruskal-Wallis test [p=0.049] with Dunn's multiple comparisons test, *$P$ < 0.05).

(B) Burst duration of the contralesional right flexor and extensor (n=11 mice, Kruskal-Wallis test [p=0.042] with Dunn's multiple comparisons test, *$P$ < 0.05).

(C) Burst duration of the ipsilesional right flexor and extensor (n=11 mice, Kruskal-Wallis test [p=0.005] with Dunn's multiple comparisons test, *$P$ < 0.05).

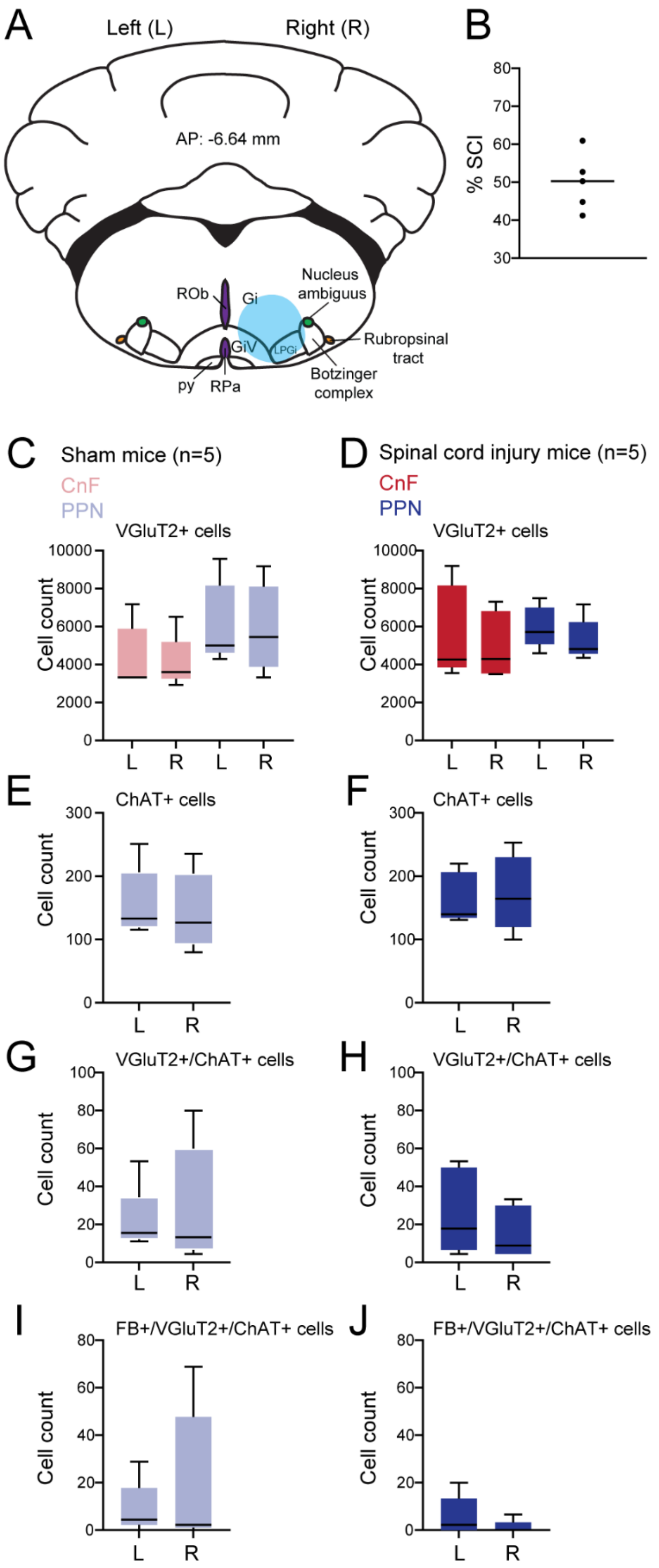

A
Left (L)
Right (R)
AP: -6.64 mm
ROb
Gi
Nucleus ambiguus
Rubropsinal tract
GiV
LPGi
Botzinger complex
py
RPa
B
% SCI
C
Sham mice (n=5)
CnF
PPN
VGluT2+ cells
Cell count
D
Spinal cord injury mice (n=5)
CnF
PPN
VGluT2+ cells
E
ChAT+ cells
F
ChAT+ cells
G
VGluT2+/ChAT+ cells
H
VGluT2+/ChAT+ cells
I
FB+/VGluT2+/ChAT+ cells
J
FB+/VGluT2+/ChAT+ cells
L
R

**Fig. S2: Cell counts of glutamatergic, cholinergic, and glutamatergic/cholinergic CnF and PPN neurons after chronic SCI.**

(A) Fast Blue injections in the gigantocellular reticular nucleus (Gi), the lateral para-gigantocellular reticular nucleus (LPGi), and the ventral pars of the gigantocellular reticular nucleus (GiV).

(B) Extent of the lesion site.

(C-D) Number of VGlut2+ neurons in left (L) ipsi- and right (R) contralesional CnF and PPN of SHAM (C) and SCI (D) mice.

(E-F) Number of ChAT+ neurons in left (L) ipsi- and right (R) contralesional PPN of SHAM (E) and SCI (F) mice.

(G-H) Number of VGlut2+/ChAT+ neurons in left (L) ipsi- and right (R) contralesional PPN of SHAM (G) and SCI (H) mice.

(I-J) Number of FastBlue+/VGlut2+/ChAT+ neurons in left (L) ipsi- and right (R) contralesional PPN of SHAM (I) and SCI (J) mice.

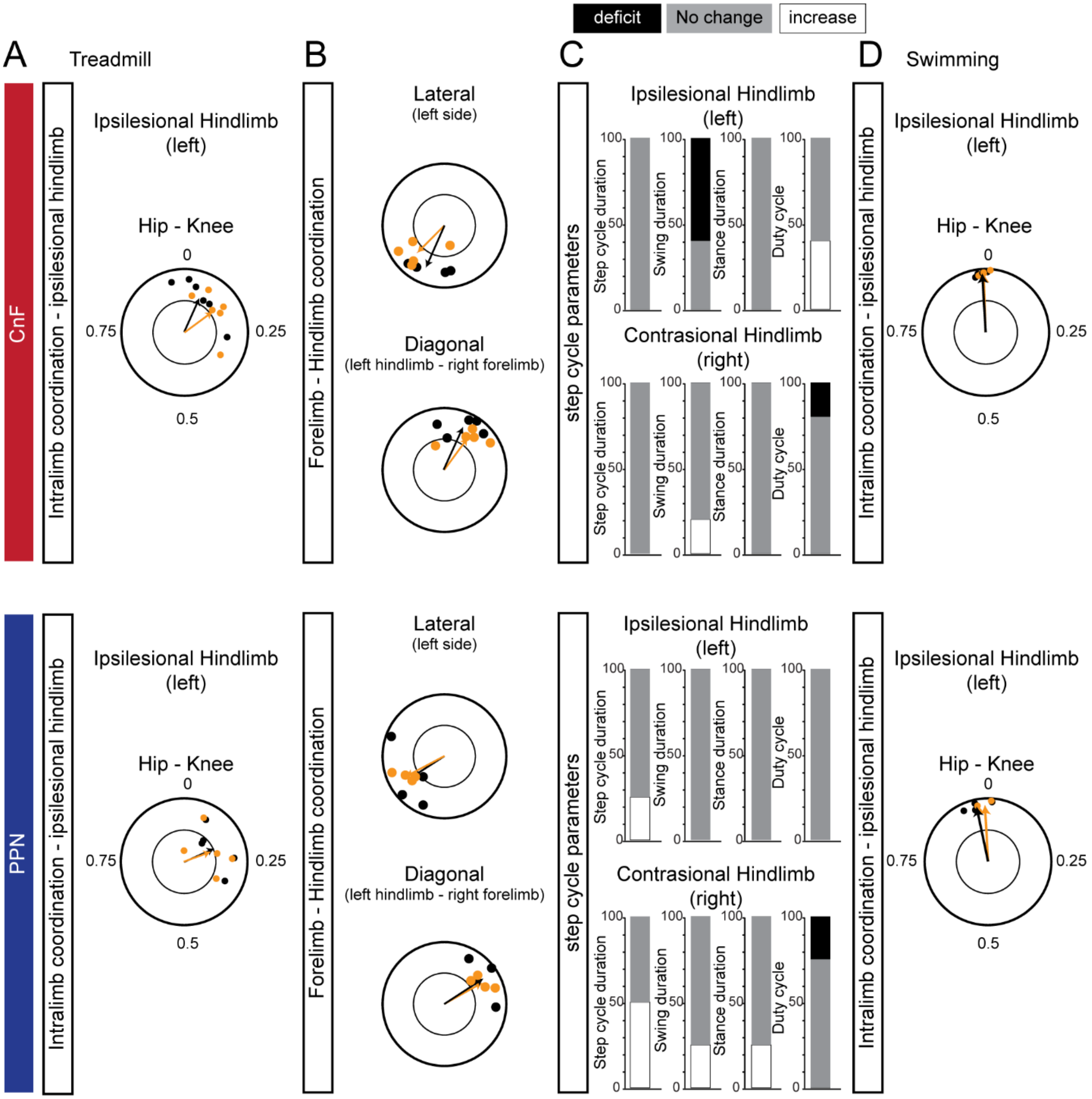


**Fig. S3: Intralimb coordination during locomotion upon genetic ablation.**

(A) Intralimb coupling between the hip and knee during treadmill locomotion before and after glutamatergic ablation of the CnF or PPN.

(B) Lateral coupling between left fore-and hindlimb (anchored on the left forelimb) and diagonal coupling between left hindlimb and right forelimb (anchored on the right forelimb).

(C) Percentage of mice exhibiting a significant deficit, improvement, or absence of change in step cycle duration, swing and stance phase duration, and duty cycle of the stance phase.

(D) Intralimb coupling between the hip and knee during swimming.

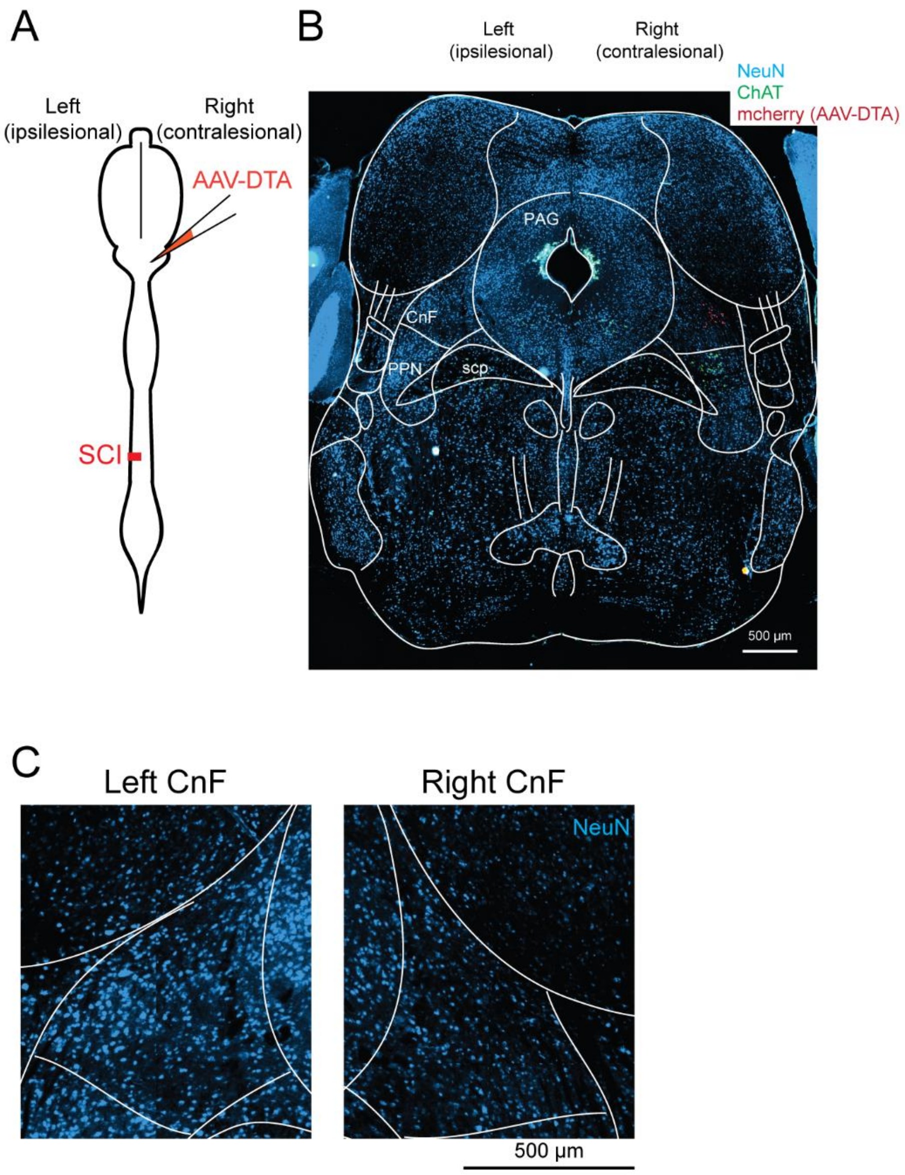


**Fig. S4: Genetic ablation of glutamatergic mesencephalic neurons.**

(A) 9 weeks after a latleral (left) thoracic hemisection, genetic ablation of glutamatergic neurons of the contralesional (right) CnF or PPN.

(B) Mcherry expression (aav-dta, red), immunostaining against chat (cholinergic neurons, green) and neun (blue) at midbrain level.

(C) High magnifications centered at the level of the left and right cnf with NeuN immunostaining.

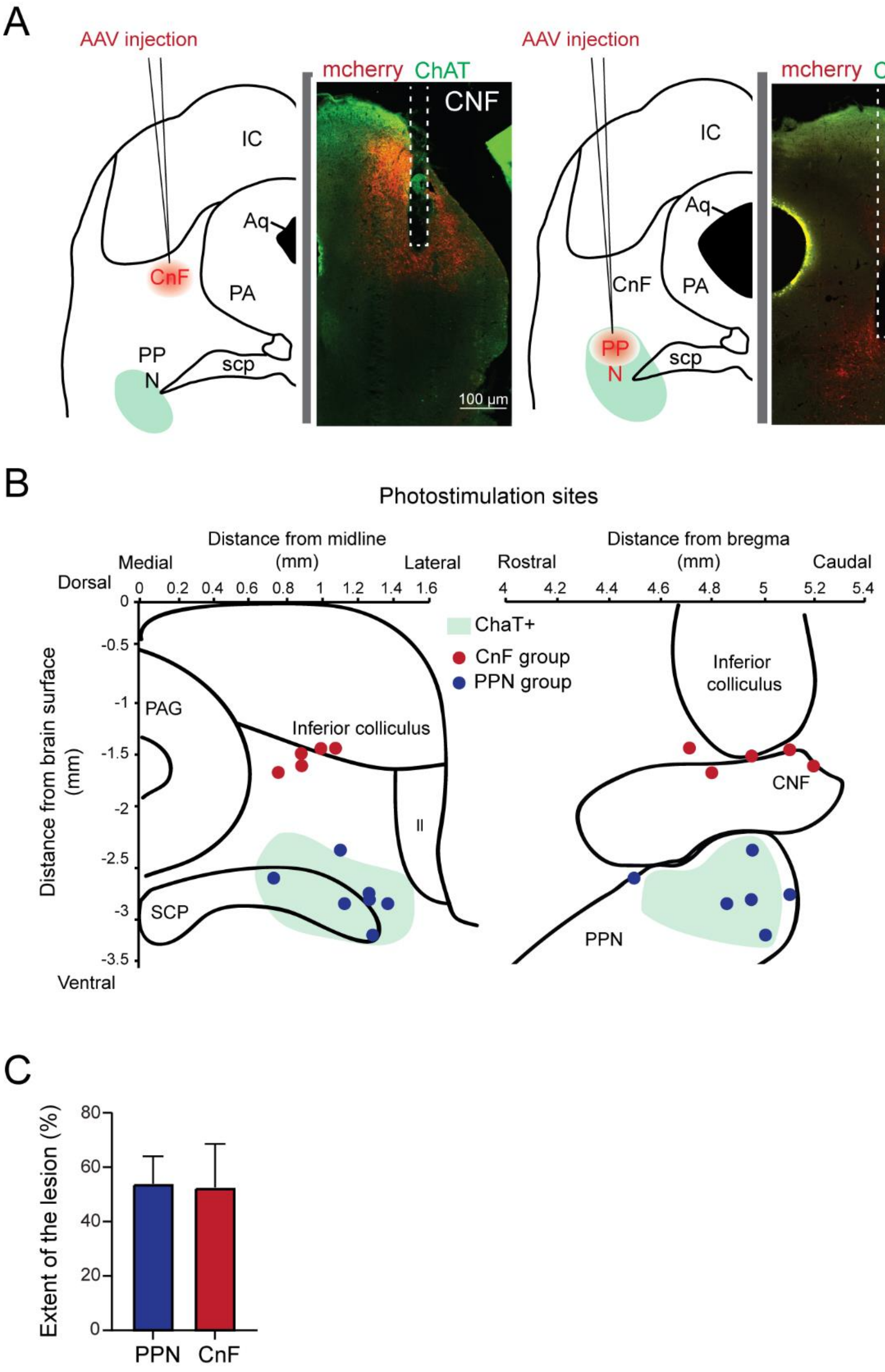


**Fig. S5: Extent of the cre-lox recombination, site of optical cannulas, and extent of the lesion site.**

(A) Extent of the cre-lox recombination.

(B) Location of the optical cannulas.

(C) Extent of the lesion site.

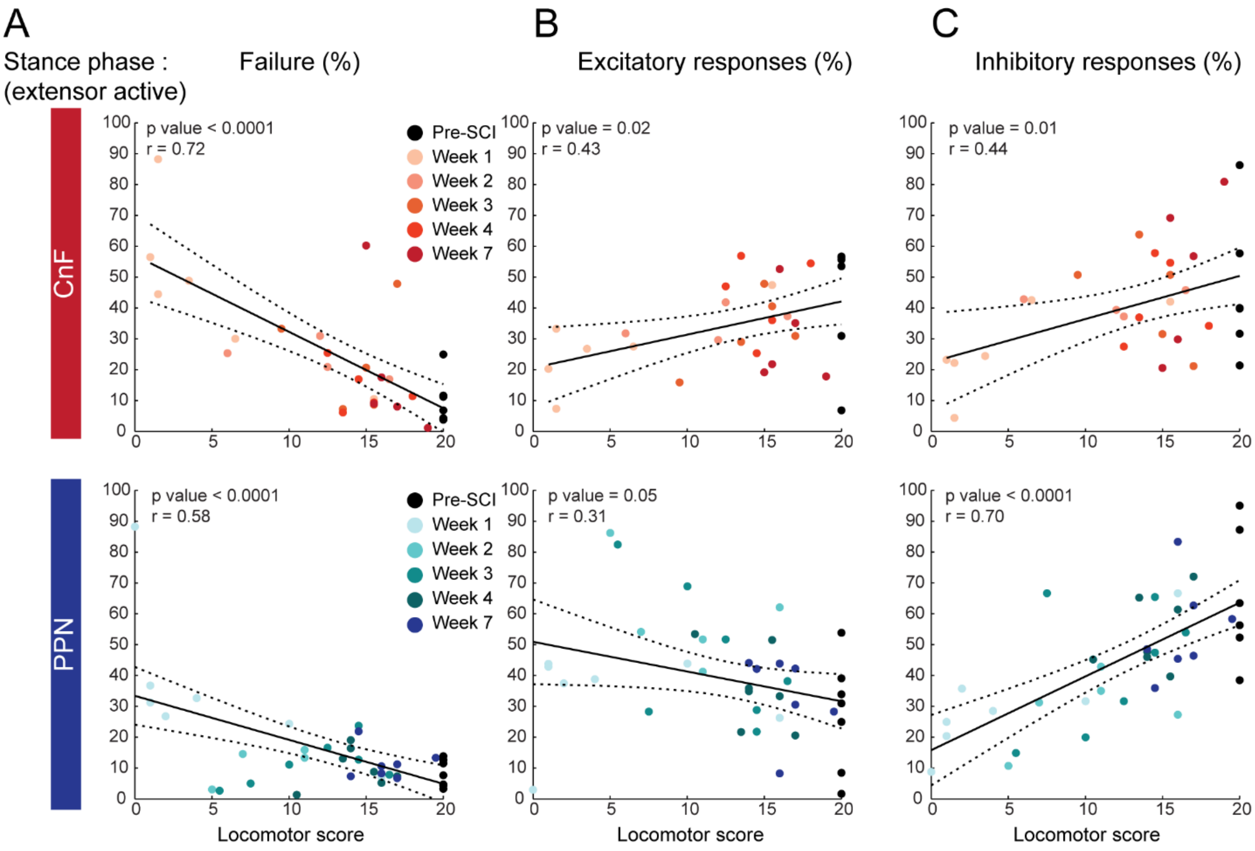


**Fig. S6: Changes in the proportion of failure, excitatory, and inhibitory motor responses in the ipsilesional extensor muscle during the stance phase after SCI.**

(A-C) Proportion of failure (A), excitatory (B), and inhibitory (C) motor responses evoked in the ipsilesional extensor during the stance phase as a function of the locomotor score of the ipsilesional hindlimb over the course of recovery after SCI.

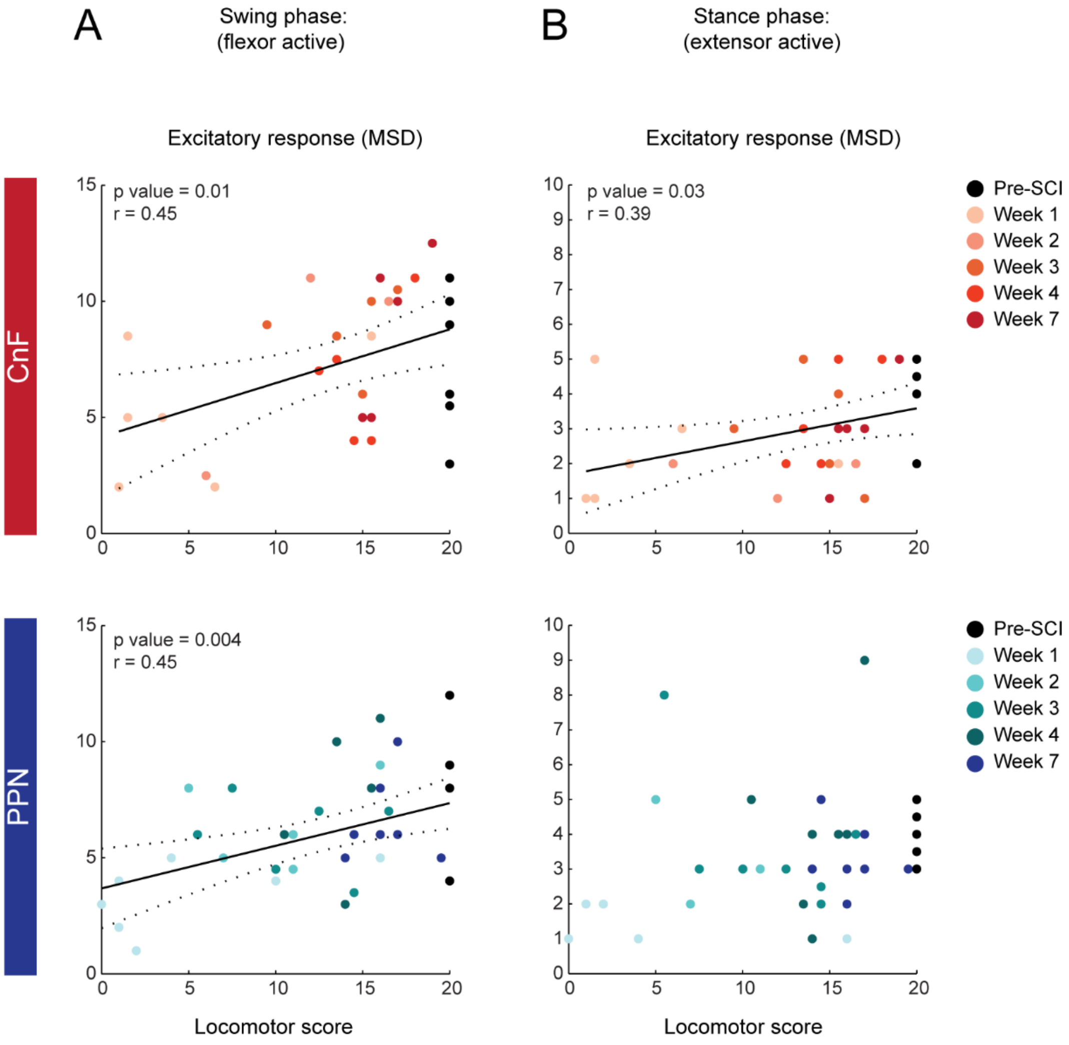


**Fig. S7: Changes in motor spike density of excitatory motor responses evoked in the ipsilesional hindlimb before and after SCI.**

(A) Motor spike density of excitatory motor responses evoked in the ipsilesional flexor muscle during the swing phase upon short pulse photo-stimulation delivered above glutamatergic CnF or PPN neurons as a function of the locomotor score of the ipsilesional hindlimb over the course of recovery after SCI.

(B) Motor spike density of excitatory motor responses evoked in the ipsilesional extensor muscle during the stance phase upon short pulse photo-stimulation delivered above glutamatergic CnF or

PPN neurons as a function of the locomotor score of the ipsilesional hindlimb over the course of recovery.

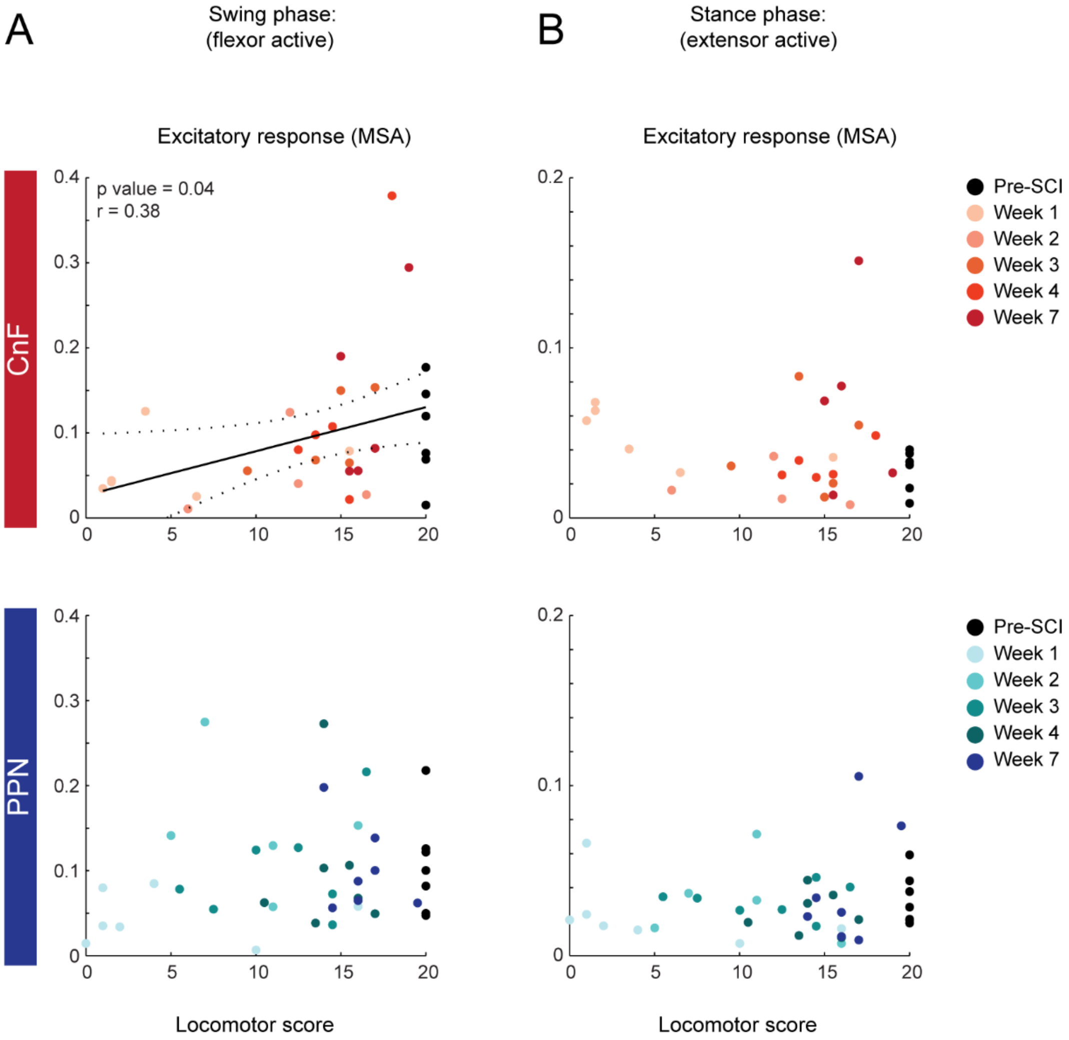


**Fig. S8: Changes in motor spike amplitude of excitatory motor responses evoked in the ipsilesional hindlimb before and after SCI.**

(A) Motor spike amplitude of excitatory responses evoked in the ipsilesional flexor muscle during the swing phase upon short-pulse photo-stimulation of glutamatergic CnF or PPN neurons as a function of locomotor score.

(B) Motor spike amplitude of excitatory responses evoked in the ipsilesional extensor muscle during the stance phase upon short-pulse photo-stimulation of glutamatergic CnF or PPN neurons as a function of locomotor score.

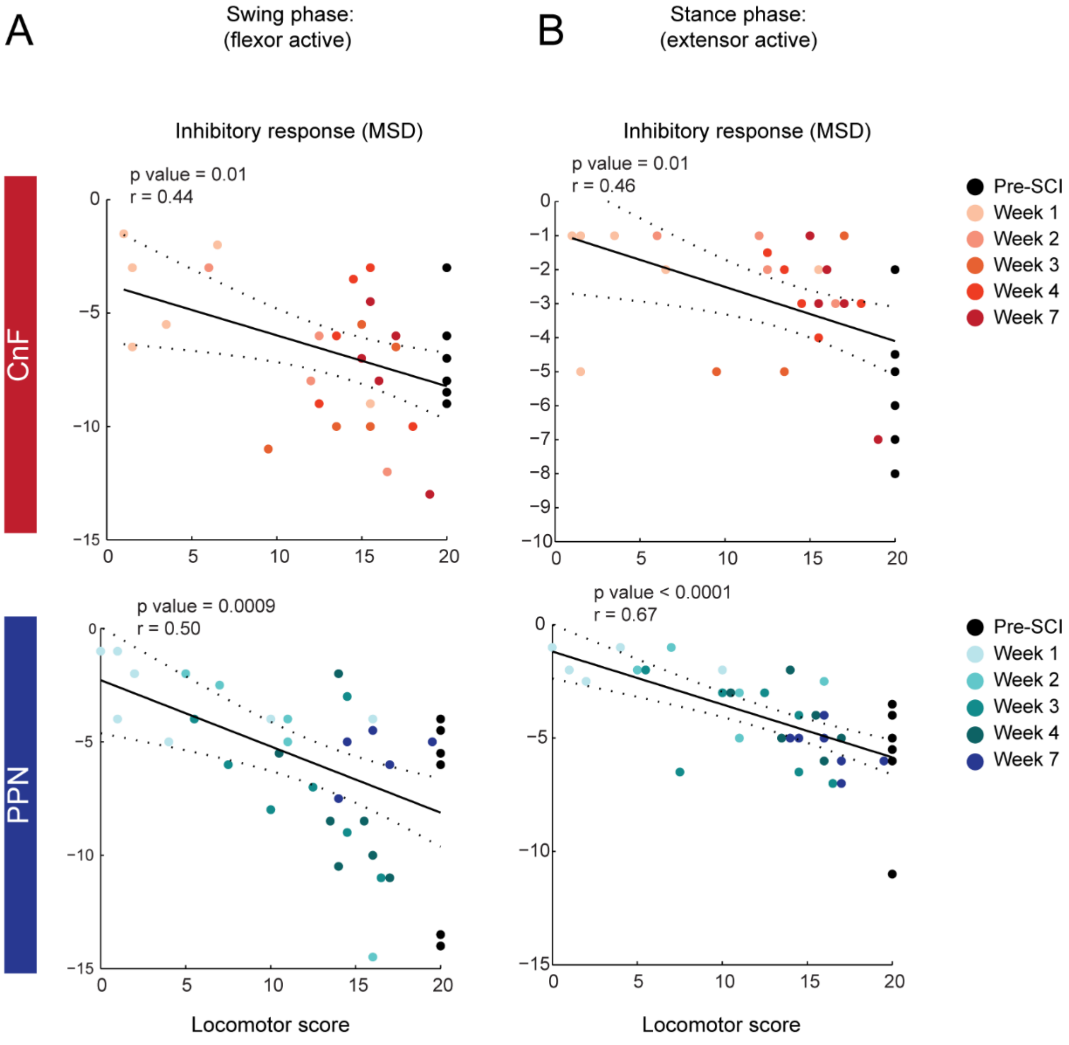


**Fig. S9: Motor spike density of inhibitory motor responses evoked in the ipsilesional hindlimb before and after SCI.**

(A) Motor spike density of inhibitory responses evoked in the ipsilesional flexor muscle during the swing phase upon short-pulse photo-stimulation of glutamatergic CnF or PPN neurons as a function of locomotor score.

(B) Motor spike density of inhibitory responses evoked in the ipsilesional extensor muscle during the stance phase upon short-pulse photo-stimulation of glutamatergic CnF or PPN neurons as a function of locomotor score.

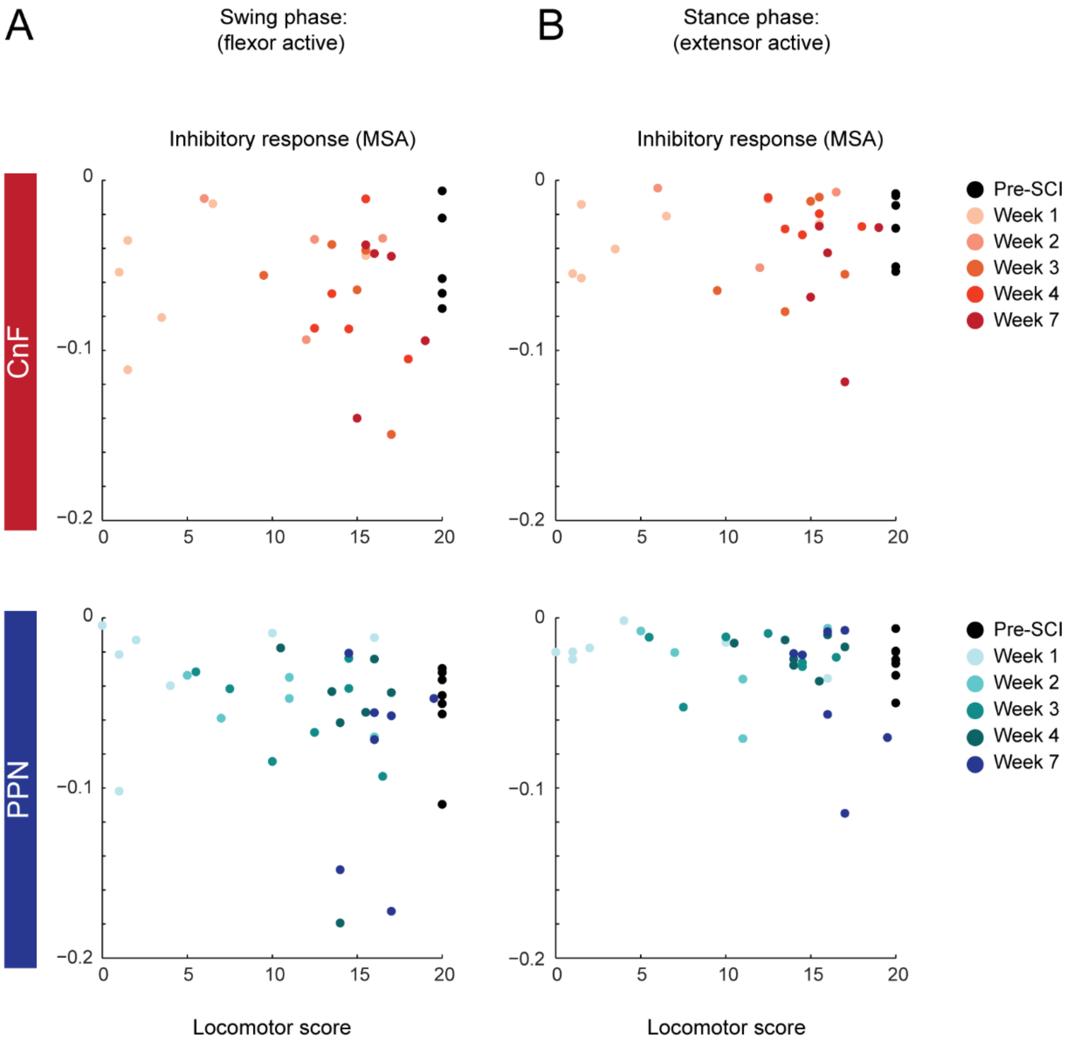


**Fig. S10: Motor spike amplitude of inhibitory motor responses evoked in the ipsilesional hindlimb before and after SCI.**

(A) Motor spike amplitude of inhibitory responses evoked in the ipsilesional flexor muscle during the swing phase upon short-pulse photo-stimulation of glutamatergic CnF or PPN neurons as a function of locomotor score.

(B) Motor spike amplitude of inhibitory responses evoked in the ipsilesional extensor muscle during the stance phase upon short-pulse photo-stimulation of glutamatergic CnF or PPN neurons as a function of locomotor score.

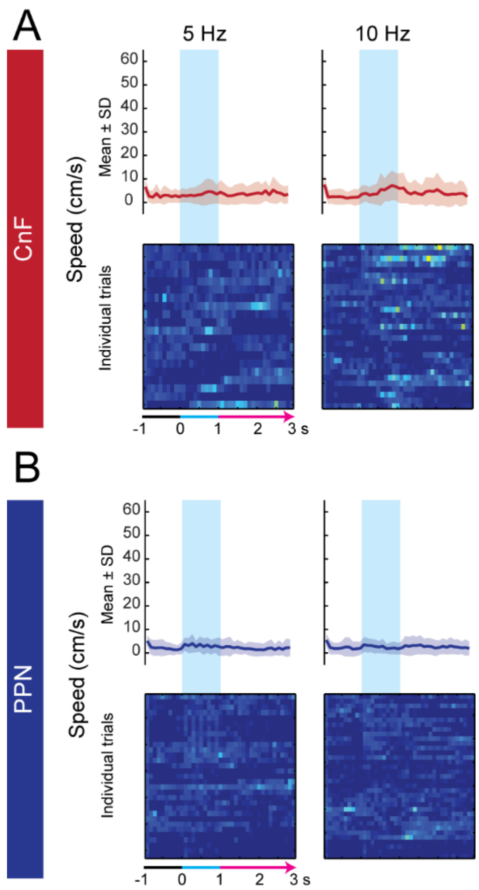


**Fig. S11: No initiation of locomotion upon photo-stimulation at 5 or 10 Hz after chronic SCI.**

(A-B) Mean and SD of locomotor speed evoked upon long train of photo-stimulation of glutamatergic neurons of the CnF (A) or PPN (B) at 5 and 10 Hz. Color-coded matrices represent individual trials (100 ms bins).

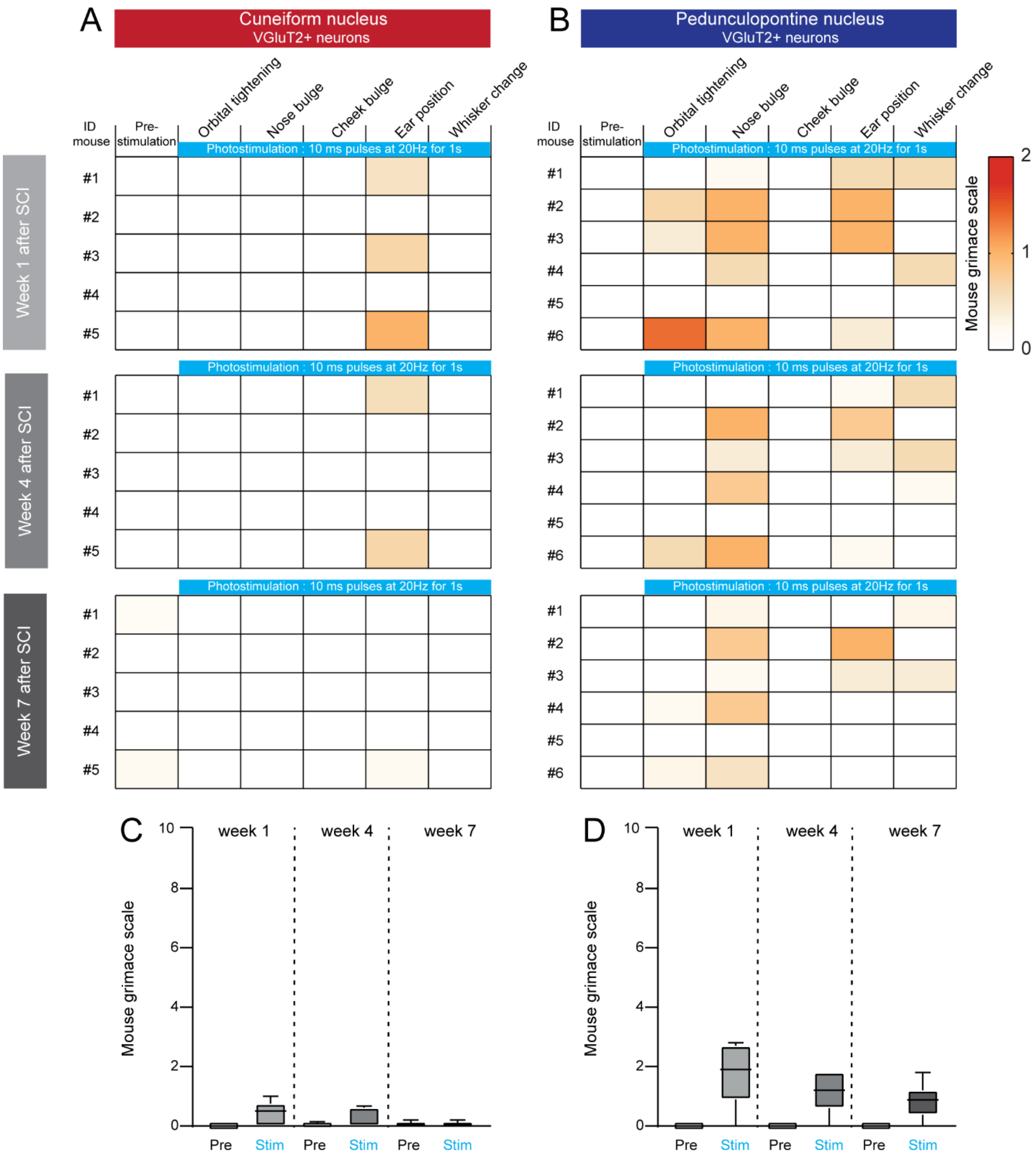


**Fig. S12: No reliable signs of stress or pain upon photo-stimulation of glutamatergic neurons of the CnF or PPN after chronic SCI.**

(A-B) Grimace scale features scored prior and during photo-stimulation of glutamatergic neurons of the CnF (A) or PPN (B) at 20 Hz at 1, 4, and 7 weeks after SCI.

(C-D) Average mouse grimace scale evoked by stimulation of the CnF (C) and PPN (D).

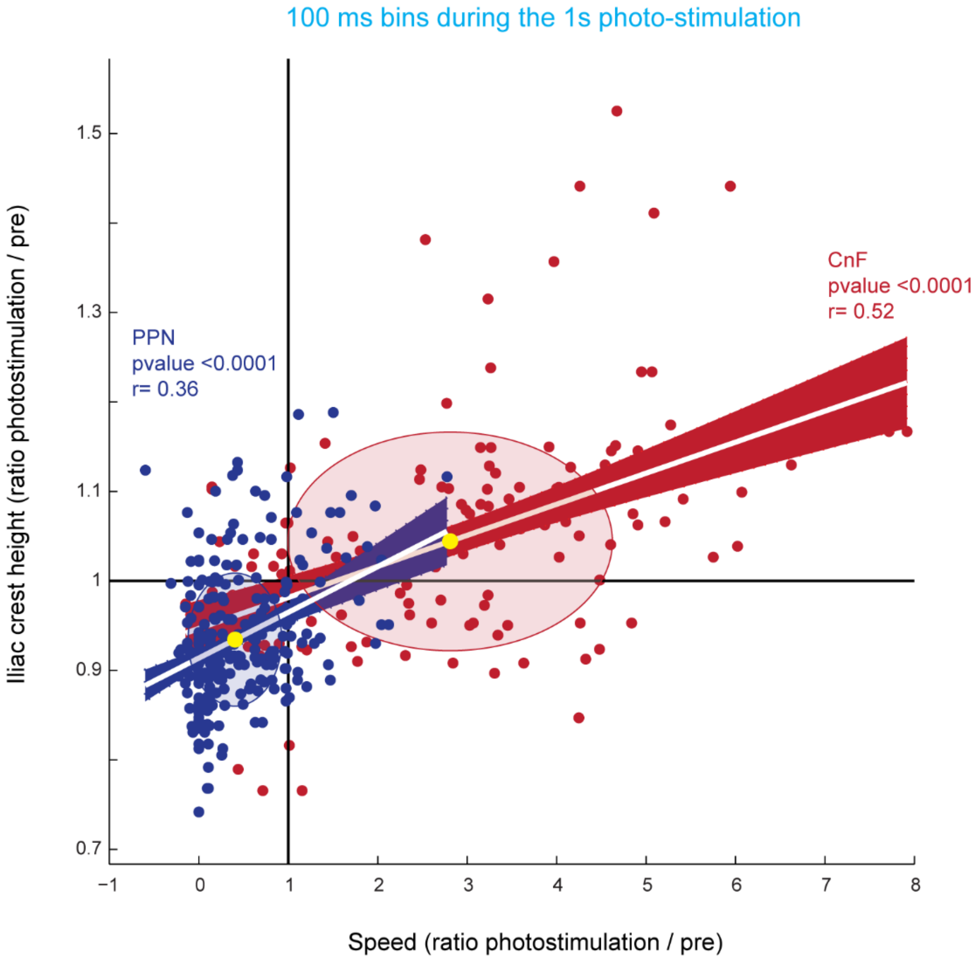


**Fig. S13: Speed and posture upon long trains of photo-stimulation of glutamatergic CnF versus PPN neurons after chronic SCI.**

Iliac crest height as a function of speed normalized on pre-stimulation data. Significant linear regressions between both parameters upon photo-stimulation of the CnF or PPN (n= 150 bins for CnF and 280 bins for PPN mice, 100 ms bins).

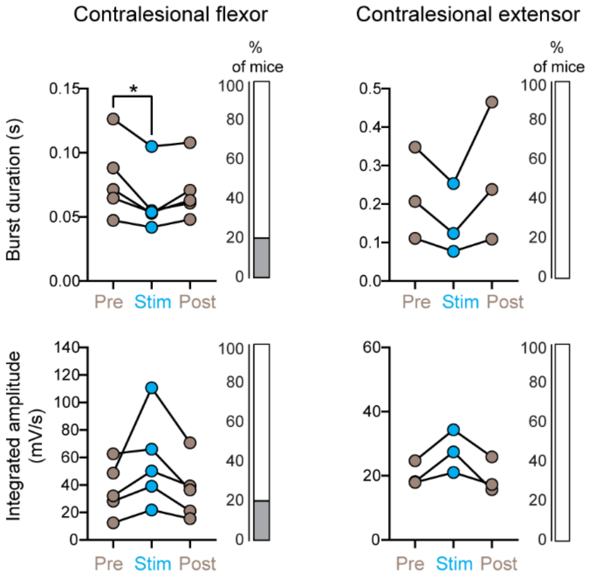


**Fig. S14: Background EMG activity of the contralesional hindlimb in response to long trains of photo-stimulation of glutamatergic CnF neurons after chronic SCI.**

Burst duration and integrated amplitude of background EMG bursts in the contralesional right flexor (left) and extensor (right). Percentage of mice showing a significant improvement (white) or no change (gray), Friedman test [p=0.009] with Dunn's multiple comparison post hoc test, **P*<0.05.

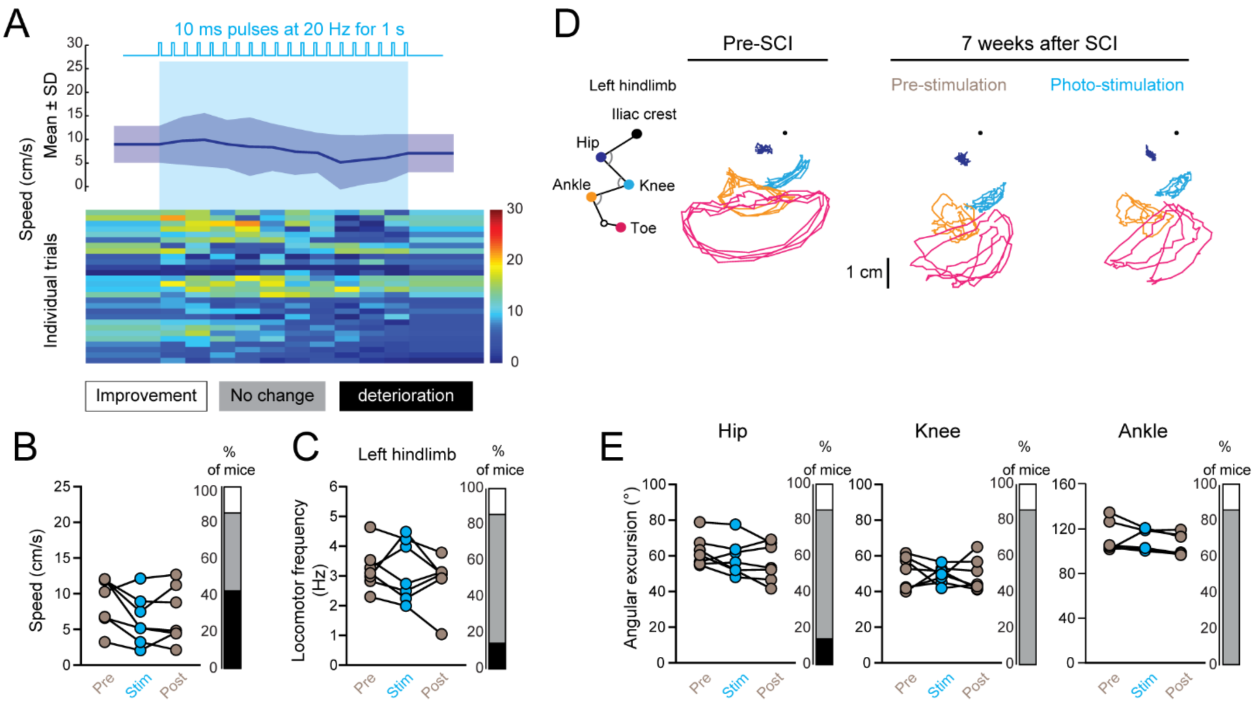


**Fig. S15: Long trains of photo-stimulation of glutamatergic PPN neurons failed to improve swimming after chronic SCI.**

(A) Mean and SD of the swimming and individual trials color-coded as a function of speed (n=7 mice, 100ms bins, 4 trials per mouse).

(B) Mean swimming speed (500 ms pre/stim/post periods, last 500 ms of the 1 s stimulation. Percentage of mice exhibiting an increase or decrease in speed upon photo-stimulation.

(C) Mean swimming frequency upon photo-stimulation. Percentage of mice exhibiting an increase or decrease in locomotor frequency upon photo-stimulation.

(D) Trajectories of the hip, knee, ankle, and toe anchored to the iliac crest during a swimming bout before and 7 weeks after SCI.

(E) Mean angular excursion of the ipsilesional hip, knee, and ankle. Percentage of mice exhibiting a significant increase, decrease, or absence of changes in their angular excursion upon photo-stimulation.

| Measured parameter | *P* | *P* | Figure |
|---|---|---|---|
| Locomotor score.<br>(left hindlimb) | $P = 0.0001$<br>Friedman test | Pre-SCI versus Week 1: $P < 0.01$<br>Dunn's multiple comparisons post hoc test | Fig 1C |
| Forward placement.<br>(left hindlimb) | $P < 0.0001$<br>Friedman test | Pre-SCI versus Week 1: $P < 0.01$<br>Dunn's multiple comparisons post hoc test | Fig 1F |
| EMG integrated amplitude.<br>(left flexor) | $P = 0.0095$<br>Kruskal-Wallis | Week 1 versus Week 4: $P < 0.05$<br>Week 1 versus Week 7: $P < 0.05$<br>Dunn's multiple comparisons post hoc test | Fig 1I |
| EMG integrated amplitude.<br>(left extensor) | $P < 0.0001$<br>Kruskal-Wallis | Week 1 versus Week 4: $P < 0.05$<br>Week 1 versus Week 7: $P < 0.05$<br>Dunn's multiple comparisons post hoc test | Fig 1J |
| Coupling between the angle of the ipsilesional hip and ankle. Pre versus after DTA. PPN group. | $P = 0.006$<br>T-test | | Fig 3E |
| Speed pre versus after DTA, swimming. | $P = 0.0006$ (CnF group)<br>Paired t-test | | Fig 3J |
| Latency to initiate full body movement, CnF group. | $P = 0.001$<br>Mann-Whitney test | | Fig 5E |
| Speed, CnF group. | $P = 0.04$<br>Friedman test | Pre versus stim: $P < 0.05$<br>Dunn's multiple comparison post hoc test | Fig 6C |
| Step frequency, CnF group. | $P = 0.0003$<br>Kruskal Wallis test | Pre versus stim: $P < 0.01$<br>Stim versus post: $P < 0.001$<br>Dunn's multiple comparison post hoc test | Fig 6C |
| Step height, CnF group. | $P = 0.009$<br>Kruskal Wallis test | Pre versus stim: $P < 0.01$<br>Dunn's multiple comparison post hoc test | Fig 6C |
| Speed, PPN group. | $P = 0.02$<br>Friedman test | Pre versus stim: $P < 0.05$<br>Dunn's multiple comparison post hoc test | Fig 6E |
| Iliac crest height, PPN group. | $P = 0.008$<br>Friedman test | Pre versus stim: $P < 0.01$<br>Dunn's multiple comparison post hoc test | Fig 6E |
| Step frequency, PPN group. | $P = 0.0001$<br>Kruskal Wallis test | Pre versus stim: $P < 0.0001$<br>Pre versus post: $P < 0.01$<br>Dunn's multiple comparison post hoc test | Fig 6E |
| EMG burst duration, left flexor. | $P = 0.009$<br>Friedman test | Pre versus stim: $P < 0.05$<br>Dunn's multiple comparison post hoc test | Fig 7E |
| EMG integrated amplitude,<br>left flexor. | $P = 0.008$<br>Repeated measures one-way ANOVA | Pre versus stim: $P < 0.05$<br>Tukey's comparison test | Fig 7E |
| Speed. | $P = 0.0004$<br>Repeated measures one-way ANOVA | Pre versus stim: $P < 0.001$<br>Pre versus post: $P < 0.05$<br>Stim versus post: $P < 0.01$<br>Tukey's multiple comparisons test | Fig 8C |
| Locomotor frequency. | $P = 0.02$<br>Friedman Test | Pre versus stim: $P < 0.05$<br>Dunn's multiple comparisons test | Fig 8D |
| Angular excursion, knee. | $P = 0.004$<br>Repeated measures one-way ANOVA | Pre versus stim: $P < 0.01$<br>Stim versus post: $P < 0.05$<br>Tukey's multiple comparisons test | Fig 8F |

**Table S1: Statistical analysis and p-values.**

Statistical tests for all main figures, normality was assessed before choosing the statistical test to perform (see material and method section).
